# Neural correlates of learning pure tones versus natural sounds in the auditory cortex

**DOI:** 10.1101/273342

**Authors:** Ido Maor, Ravid Shwartz-Ziv, Libi Feigin, Yishai Elyada, Haim Sompolinsky, Adi Mizrahi

## Abstract

Auditory perceptual learning of pure tones causes tonotopic map expansion in the primary auditory cortex (A1), but the function this plasticity sub-serves is unclear. We developed an automated training platform called the ‘Educage’, which was used to train mice on a go/no-go auditory discrimination task to their perceptual limits, for difficult discriminations among pure tones or natural sounds. Spiking responses of excitatory and inhibitory L2/3 neurons in mouse A1 revealed learning-induced overrepresentation of the learned frequencies, in accordance with previous literature. Using a novel computational model to study auditory tuning curves we show that overrepresentation does not necessarily improve discrimination performance of the network to the learned tones. In contrast, perceptual learning of natural sounds induced ‘sparsening’ and decorrelation of the neural response, and consequently improving discrimination of these complex sounds. The signature of plasticity in A1 highlights its central role in coding natural sounds as compared to pure tones.

## INTRODUCTION

Learning is accompanied by plastic changes in brain circuits. This plasticity is often viewed as substrate for improving computations that sub-serve learning and behavior. A well-studied example of learning-induced plasticity is following perceptual learning where cortical representations change towards the learned stimuli ^1,2^. Whether such changes improve discrimination is still debated and the mechanisms leading to these changes are still largely unknown. Furthermore, learning paradigms in animal models are often limited to simplified, unnatural stimuli that deviate further from real learning experience and that brain circuits actually compute.

Perceptual learning is an implicit form of lifelong learning during which perceptual performance improves with practice ^3^. Perceptual learning spans a wide range of sensory modalities and tasks like learning and improving reading skills, acquiring language, or the learning to discriminate different shades of color and flavors. Extensive psychophysical research on perceptual learning tasks led to a general agreement on some attributes of this type of learning ^4^. For example, perceptual learning has been shown to be task specific, poorly generalized to other senses or tasks. It is also largely agreed upon that gradual training is essential for improvement ^5–14^ Given the specificity observed at the behavioral level, functional correlates of perceptual learning are thought to involve neural circuits as early as primary sensory regions ^2,15^. In auditory learning paradigms, changes are already observed at the level of primary auditory cortex ^16^. Learning to discriminate among tones results in tonotopic map plasticity towards the trained stimulus ^17–20^; reviewed in ^21^. Notably, however, not all studies could replicate the learning induced changes in the tonotopic map ^22^. Furthermore, artificially induced map plasticity was shown to be unnecessary for better discrimination performances *per se*, ^23,24^. Our understanding of the mechanisms underlying auditory cortex plasticity remains rudimentary, let alone for more natural stimuli.

To gain understanding of learning-induced plasticity at single neuron resolution, animal models have proven very useful. Mice, for example, offer the advantage of a rich genetic experimental toolkit to study neurons and circuits with high efficiency and specificity ^25^. Historically, the weak aspect of using mice as a model was its limited behavioral repertoire to learn difficult tasks. However, in the past decade technical difficulties to train mice to their limits have been steadily improving with increasing number of software and hardware tools to probe mouse behavior in high resolution ^26–33^. Here, we developed our own experimental system for training groups of mice on an auditory perceptual task – an automatic system called the ‘Educage’. The Educage is a simple affordable system that allows efficient training of several mice simultaneously to discriminate among pure tones or complex sounds.

A1 is well known for its tonotopic map plasticity following simple forms of learning in other animal models ^21^. An additional interest in primary auditory cortex is its increasing recognition as a brain region involved in coding complex sounds ^34,35^. We thus asked what are the similarities and differences in the plastic changes single neurons undergo following training to discriminate pure tones or natural stimuli. We describe distinct changes in the long-term stimulus representations by L2/3 neurons of mice following perceptual learning and assess how these contribute to information processing by local circuits. Using two photon targeted electrophysiology, we also describe how L2/3 parvalbumin-positive neurons change with respect to their excitatory counterparts. Our work provides a behavioral, physiological and computational foundation to questions of auditory-driven plasticity in mice.

## METHODS

### Animals

A total of n=88, 10-11 week-old female mice were used in this work as follows. 44 mice were C57BL/6 mice and 44 mice were PV–Cre; Ai9 double-heterozygous (PV × Ai9;^36,37^). All experiments were approved by the Hebrew University’s IACUC.

### Behavioral setup

The ‘Educage’ is a small chamber (10×10×10 cm), opening on one end into a standard animal home cage where mice can enter and exit freely (Fig. 1a and Supplementary figure 1a). On the other end, the chamber contains the hardware that drives the system, hardware for identifying mice and measuring behavioral performance. Specifically, at the port entrance there is a coil radio antenna (ANTC40 which connected to LID665 stationary decoder; Dorset) followed by infra-red diodes used to identify mice individually and monitor their presence in the port. This port is the only access to water for the mice. Water is delivered *via* a solenoid valve (VDW; SMC) allowing precise control of the water volume provided on each visit. Water is delivered via a water spout, which is also a lickometer (based on 1 microampere current). An additional solenoid valve is positioned next to the water spout in order to deliver a mild air puff as a negative reinforcement, if necessary. For sound stimulation, we positioned a speaker (ES1; TDT), at the center of the top wall of the chamber. Sound was delivered to the speaker at 125 kHz sampling rate *via* a driver and a programmable attenuator (ED1, PA5; TDT). For high speed data acquisition, reliable control, on-board sound delivery, and future flexibility, the system was designed via a field programmable gate array (FPGA) module and a real-time operating system (RTOS) module (CompactRIO controller; National Instruments). A custom-made A/D converter was connected to the CompactRIO controller, mediated signal from infra-red diodes and lickometer and controlled the valves. A custom code was written in Labview to allow an executable user-friendly interface with numerous options for user-input and flexibility for designing custom protocols. All software and hardware design are freely available for download at https://github.com/MizrahiTeam/Educage.

**Figure 1-.**
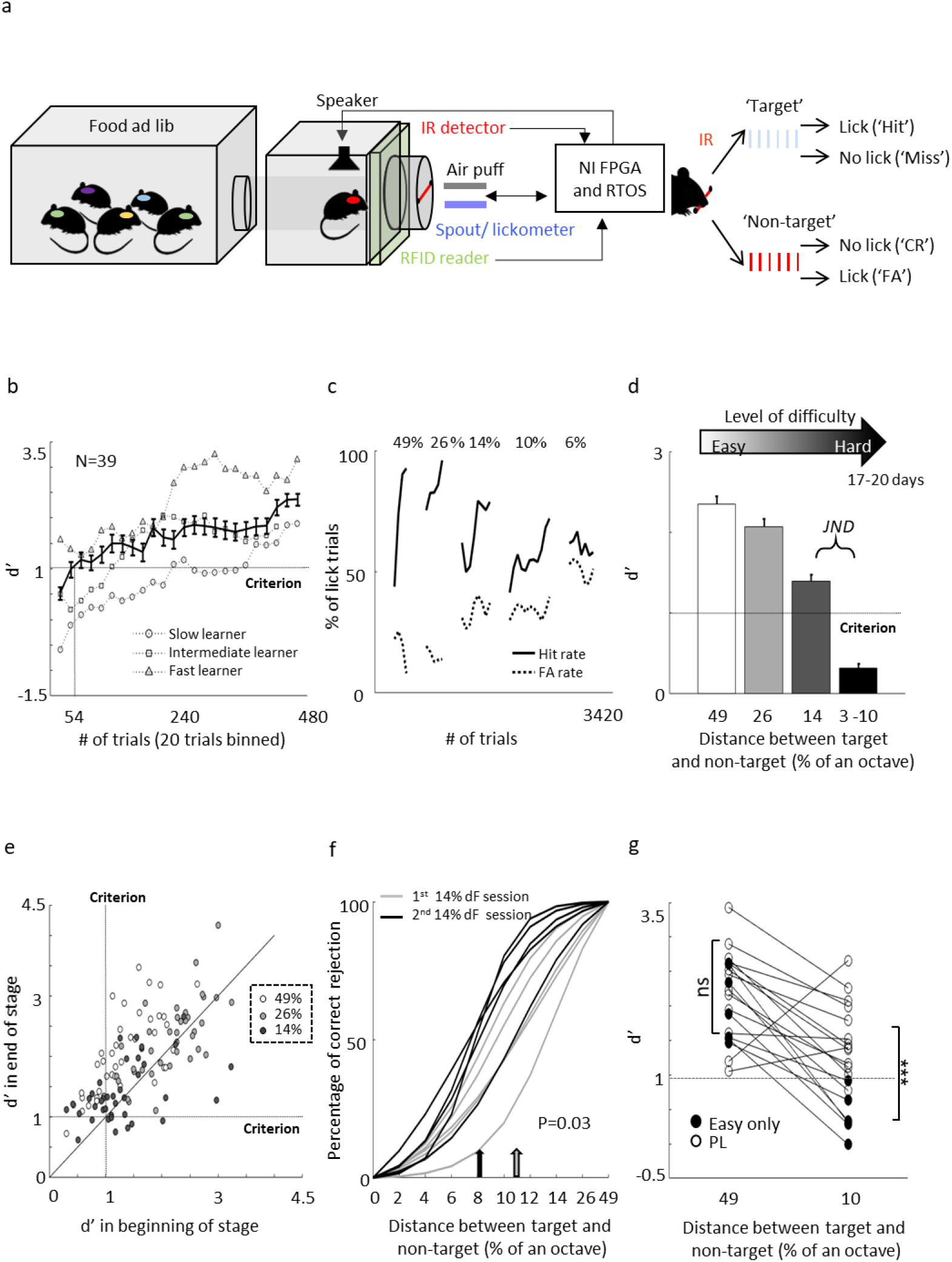
Perceptual learning in the ‘Educage’. **A. Left:** Schematic design of the ‘Educage’ system and its components. **Right:** Schematic representation of the go/no-go auditory discrimination task. CR-correct rejection, FA-False Alarm. **B.** d’ values of three representative mice and the population average± s.e.m for the first stage of discrimination. Learning criterion is represented as dashed line (d’=1). **C.** Lick responses to the target tone (solid line) and non-target tone (dashed line) of one representative mouse along different discrimination stages. **D.** population average d’ values for the different discrimination stages. N=39 mice (mean ± s.e.m). Shades denote the level of difficulty (from 49% to 4-10% octave apart). **E.** Individual d’ values at the end of each level as a function of d’ in the beginning of that level. Shades denote the level of difficulty. Learning criterion represented as dashed lines (d’=1). **F.** Normalized psychometric curves of five mice calculated from the first (light curves) and the second (dark curves) 14%/octave session. Light and dark arrows indicate average decision boundaries in the first and second sessions respectively (Mann-Whitney U-test on criteria: p=0.03). **G.** d’ values in easy (49%/octave) and more difficult (10% /octave) discrimination stages of individual mice from the ‘Easy only’ group (filled circles) and from the perceptual learning group (blank circles). d’ values are significantly different between groups only for the hard discrimination level (Mann-Whitney U-test: ***p<0.001).

### Training paradigm

Prior to the training, each mouse was implanted, under light and very short period of Isoflurane anesthesia, with Radio Frequency Identification (RFID; Trovan) chip on the back of its head. RFID chips allow identification of mice individually, which is then used by the system to control the precise stimulus delivery and track behavioral performance, on a per-mouse basis. Food and water were provided *ad libitum*. While access to water was only in the Educage, mice could engage the water port without restriction. Thus, mice were never deprived of food nor water. At the beginning of each experiment, RFID-tagged mice were placed in a large home-cage that was connected to the Educage. Before training, we removed the standard water bottle from the home cage. Mice were free to explore the Educage and drink the water at the behavioral port. Every time a mouse entered the behavioral port it was identified individually by the antenna and the infra-red beam and a trial was initiated. Before learning, any entry to the port immediately resulted in a drop of water, but no sound was played. Following 24hrs of exploration and drinking, mice were introduced for the first time to the ‘target’ stimulus – a series of six 10 kHz pure tones (100 ms duration, 300 ms interval; 3 ms on and off linear ramps; 62 dB SPL) played every time the mouse crossed the IR beam. To be rewarded with water, mice were now required to lick the spout during the 900 ms response window. This stage of operant conditioning lasted for 2-3 days until mice performed at >70% hit rates.

We then switched the system to the first level of discrimination when mice learned to identify a non-target stimulus (a 7.1 kHz pure tone series) from the already known target stimulus (separated by 49%/octave). Thus, on each trial one of two possible stimuli were played for 2.1 seconds-either a target tone or a non-target tone. Mice were free to lick the water spout anytime, but for the purpose of evaluating mouse decisions we defined a response window and counted the licking responses only within 900 ms after the sound terminated. Target tones were played at 70% probability and non-target tones at 30% probability, in a pseudorandom sequence. A ‘lick’ response to a target was rewarded with a water drop (15 microliter) and considered as a ‘Hit’ trial. A ‘no lick’ response to a target sound was considered as a ‘Miss’ trial. A lick response to the nontarget was considered a ‘False Alarm’ (FA) trial, which was negatively reinforced by a mild air puff (60 PSI; 600ms) followed by a 9 second ‘timeout’. A ‘no lick’ response to the non-target was considered a correct rejection (CR) and was not rewarded.

Once mice learned the easy discrimination, we switched the system to the second discrimination stage. Here, we increased task difficulty by changing the non-target tone to 8.333 kHz, thus decreasing the inter tone distance to 26%/octave. Then, at the following stage, the inter tone distance was further decreased to 14%/octave and then down to 6%/octave. This last transition was often done in a gradual manner (»12%/octave»10%/octave»8%/octave»6%/octave). In some of the animals (n=25), we trained mice to their JND and then changed the task back to an easier level. For some of the mice (n=13) we played ‘catch trials’ during the first and second sessions of the 14%/octave discrimination stages. In catch trials, different tones spanning the frequency range of the whole training (7-10 kHz) were presented to the animals in low probability (6% of the total number of sounds), and were neither negatively nor positively reinforced.

For the vocalizations task, we used playback of pups’ wriggling calls (WC) as the target stimulus. These vocalizations were recorded with a one-quarter inch microphone (Brüel & Kjær) from P4–P5 PV × Ai9 pups (n = 3), were sampled at 500 kHz and identified offline (Digidata 1322A; Molecular Devices). As the non-target stimulus, we used manipulations of the WC. During the first stage of the operant learning, mice learned to discriminate between WC and a fully reversed version of this call. Then, the second manipulation on the non-target stimulus was a gradual change of the frequency modulation (FM) of each syllable in the call while leaving the temporal structure of the call intact. To manipulate the syllable FM we used a dynamic linear FM ramp. This operation multiplies each sampling interval within the syllable, by a dynamic speeding factor, which changed according to the relative distance from the start and end of the syllable, and generated a new waveform by interpolation from the original waveform. For example, for a 0.6 speeding factor, the beginning of each syllable was slower by a factor of 0.4 while the end of each syllable accelerated by a factor of 0.4. The range of sound modulation used here was 0.66-0.9. A value of 0.66 is away from the WC, 0.9 similar to the WC and 1 exactly the same as the WC. The basic task design for the non-target sound was as follows: Reverse» 0.66 »0.81» 0.9.

### Surgical procedure

Mice were anesthetized with an intraperitoneal injection of ketamine and medetomidine (0.80 and 0.65 mg/kg, respectively) and a subcutaneous injection of Carprofen (0.004 mg/g). Additionally, dextrose–saline was injected to prevent dehydration. Experiments lasted up to 8 h. The depth of anesthesia was assessed by monitoring the pinch withdrawal reflex, and ketamine/medetomidine was added to maintain it. The animal’s rectal temperature was monitored continuously and maintained at 36 ± 1°C. For imaging and recording, a custom-made metal pin was glued to the skull using dental cement and connected to a custom stage to allow precise positioning of the head relative to the speaker (facing the right ear). The muscle overlying the left auditory cortex was removed, and a craniotomy (~2 × 2 mm) was performed over A1 (coordinates, 2.3 mm posterior and 4.2 mm lateral to bregma) as described previously ^38–40^.

### Imaging and electrophysiology

Cell-attached recordings were obtained using targeted patch-clamp recording by a previously described procedure ^40–42^. For visualization, the electrode was filled with a green fluorescent dye (Alexa Flour-488; 50 μM). Imaging of A1 was performed using an Ultima two-photon microscope from Prairie Technologies equipped with a 16X water-immersion objective lens (0.8 numerical aperture; CF175; Nikon). Two-photon excitation of the electrode and somata was used at 930 nm (DeepSee femtosecond laser; Spectraphysics). The recording depths of cell somata were restricted to subpial depths of 180–420 μm, documented by the multiphoton imaging. Spike waveform analysis was performed on all recorded cells, verifying that tdTomato+ cells in L2/3 had faster/narrower spikes relative to tdTomato-negative (tdTomato–) cells (See also^43^).

### Auditory stimuli

The auditory protocol comprised 18/24 pure tones (100 ms duration, 3 ms on and off linear ramps) logarithmically spaced between 3-40 kHz and presented at four sound pressure levels (72–42 dB SPLs). Each stimulus/intensity combination was presented 10/12 times at a rate of 1.4 Hz. The vocalizations protocol comprised the playback of pups’ wriggling calls (WC) and 3 additional frequency modulated (FM) calls, presented at 62 dB SPL for 16 repetitions.

### Behavioral data analysis

To evaluate behavioral performance we calculated, for different time bins (normally 20 trials), Hit and FA rates, which are the probability to lick in response to the target and non-target tones, respectively. In order to compensate for the individual bias, we used a measure of discriminability from signal detection theory – d-prime (d’). D’ is defined as the difference between the normal inverse cumulative distribution of the ‘hit’ and FA rates ^44^. D’ for each discrimination stage was calculated based on trials from the last 33% of the indicated stage. Psychometric curves were extracted based on mouse performance in response to the catch trials. By fitting a sigmoidal function to these curves we calculated decision boundaries as the inflection point of each curve. Detection time was calculated for each mouse individually, by determining the time in which lick patterns in the correct reject *vs*. the hit trials diverged (i.e. the time when significance levels crossed P<0.001 in a two-sample t-test).

### Data analysis - electrophysiology

Data analysis and statistics were performed using custom-written code in MATLAB (MathWorks). Spikes were extracted from raw voltage traces by thresholding. Spike times were then assigned to the local peaks of suprathreshold segments and rounded to the nearest millisecond. For each cell, we obtained peri-stimulus time histogram (PSTH) and determined the response window as the 100 ms following stimulus onset which evoked the maximal response integral. Only neurons that had tone-evoked response (determined by a two sample t-test) were included in our dataset. Based on this response window, we extracted the cell’s tuning curve and frequency-response area (FRA). Evoked firing rate was calculated as the average response to all frequencies that evoked a significant response. Firing rate in the training band was calculated as the response to frequencies inside the training band (7-10 kHz), averaged across all intensities. Best frequency (BF) of each cell was determined as the tone frequency that elicited the strongest response averaged across all intensities. The selectivity of the cell is the % of all frequency-intensity combinations that evoked significant response (determined by a two sample t-test). Pairwise signal correlations (r_sc_) were calculated as Pearson correlation between FRA’s matrices of neighboring cells. The spontaneous firing rate of the cell was calculated based on the 100 ms preceding each stimulus presentation. Response latency is the time point after stimulus onset at which average spike count reached maximum.

### Statistical model based on the Independent Basis Functions (IBF) Method

Since the measured responses before and after learning are not from the same cells, we cannot estimate the changes of individual tuning curves due to learning. Instead, we must rely on estimated learning-induced changes in the *ensemble* of single-neuron responses. Our goal, therefore, was to build a statistical model of single-neuron tuning curves before and after learning. This model is different before and after learning, and was used to estimate the learning-induced changes in the population of A1 responses. Furthermore, we use this model as a generative model that allowed us to generate a large number of ‘model neurons’ with statistically similar response properties as the measured ones.

In principle, one could use a parametric model, by fitting each observed tuning curve to a specific shape of functions (e.g., Gaussian tuning curves). However, since the tuning curves of neurons to tone frequencies do not have symmetric ‘gaussian’ shapes, and some are bimodal, fitting them to a parametric model has not been successful. Instead, we chose to model each single neuron response as a weighted sum of a small set of orthogonal basis functions,

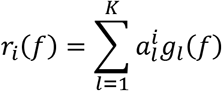

Here, *r_i_*(*f*) is the firing rate (i.e., the trial-averaged spike count) of the i-th neuron in response to the stimulus with frequency *f*; *K* is the number of orthogonal basis functions denoted by *g_l_*(*f*) (dependencies in *f* are in log scale). In order to determine the basis functions and the coefficients, 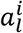 we performed Singular Value Decomposition (SVD) of the matrix of the measured neuronal firing rates for the 18 values of *f* Our model (1) uses a subset of the K modes with the largest singular values (the determination of K is described below). The SVD yields the coefficients 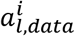 for the *N observed* neurons and (2) smoothes the resultant SVD f-dependent vectors using a simple ‘moving average’ technique to generate the basis functions, *g_l_*(*f*) (3) Importantly, to use the SVD as a generative model, for each *l* we compute the histogram of the 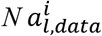 coefficients. To generate ‘new neurons’, we sample each coefficient independently from the corresponding histogram. In other words, we approximate the joint distribution of the coefficients by a factorized distribution. This allowed us to explore the effect of changing the number of neurons that downstream decoders use in order to perform the perceptual task.

Model (1) describes the variability of the population responses to the stimulus, in terms of tuning curves of the trial averages firing rates. Additional variability in the data is the single trial spike count. We model these as independent Poisson random variables with means given by *r_i_*(*f*). Since neurons are not simultaneously recorded, we do not include noise correlations in the model. We performed this procedure for the *naïve* and *expert* measured responses separately, so that both the basis functions and the coefficient histograms are evaluated for the two conditions separately. Note we do not make Gaussian assumptions about the coefficient histograms. In fact, the observed histograms are in general far from Gaussian.

### The choice of number of basis functions

Due to a limited number of trials that we sampled for each neuron, taking a large value of *K* can result in over-fitting the model to the noise caused by the finite number of trials. To estimate the optimal number of basis functions, we evaluated the percentage of response firing rate variability of the population (i.e, the fraction of the sum of the squared SVD eigenvalues) as a function of K. We also evaluated the parameters of model (1) based on a subset of trials and checked how well it accounts for the observed tuning curves that are calculated from the test trials. We took *K* that produces the smallest test error and saturated the fractional variance.

We used Model (1) with the above choice of *K* in order to evaluate the discrimination ability of the population of A1 neurons, by creating *an ensemble* of single-neuron responses for the naïve and expert conditions. To generate the model neurons, we sampled the coefficients of the basis function independently from the corresponding histogram of the measured neurons, and used these neurons for the calculations depicted below.

### Fisher Information

We calculated the Fisher Information (FI) for each condition (naïve vs. expert) using our model (1). FI measure bounds the mean squared error of an (unbiased) estimator of the stimulus from the noisy single trial neuronal responses. When the neuronal population is large (and they are noise-independent) FI also determines the discriminability d’ of a maximum likelihood discriminator between two nearby values of the stimulus ^45^. Under the above Poisson assumption, the FI for the i-th neuron is equal to 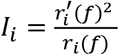, where 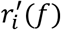 is the derivative of the firing rate with respect to the stimulus value *f*. The total FI is the sum of the FIs of individual neurons ^45^. This has been evaluated in both naïve and expert conditions. Note that the FI are functions of the stimulus value f around which the discrimination task is performed.

### Discrimination by linear readout

We applied a linear decoder to assess the ability to discriminate between nearby stimuli on the basis of the neuronal population responses. We trained a support vector machine (SVM) with a linear kernel, which finds an ‘optimal’ linear classifier that discriminate between two nearby frequencies on the basis of single-trial vectors of spike counts generated with our generative model (1) and Poisson variability. We then evaluated the probability of classification errors to test trials, in both naïve and expert conditions. Since our training set is not linearly separable, we used SVM with slack variables ^46^, which incorporates a ‘soft’ cost for classification errors. Each classification was iterated maximum 500 times (or until converged). In each iteration, 16 trials were used for training the classifier and 4 trials were used to test the decoder accuracy.

### Data analysis – vocalization responses

Similarity of response to different vocalizations was calculated as Pearson correlation between the PSTH’s of the different stimuli. To quantify life time sparseness we used the following measure S= (1 − [(Σr_i_/n)2/Σ(ri 2/n)])/[1 − (1/n)], where r_i_ is the response to the ith syllable in the original vocalization (averaged across trials) and n is the number of syllables. Values of S near 0% indicate a dense code, and values near 100% indicate a sparse code ^47^. Population sparseness was calculated as 100-the percent of cells that evoked a significant response to each syllable in the call ^48^. Classification of vocalization identity based on population activity was determined using the Support Vector Machine (SVM) decoder with a linear kernel and slack variables. The decoder was tested for its accuracy to differentiate between responses to two different vocalizations. The input to the SVM consisted of the spike count of each neuron in the syllable response window. The same number of neurons (37) was used in both groups to avoid biases. We then evaluated the probability of classification errors to test trials, using leave-one-out cross validation. Each classification was iterated 1000 times. In each iteration, 15 trials were used for training the classifier and one trial was used to test decoder accuracy. The number of syllables utilized in the decoder were increased cumulatively.

## RESULTS

### Behavior – Discrimination of pure tones

To study perceptual learning in mice, we developed a behavioral platform named the ‘Educage’ (all software and hardware design are freely available for download at https://github.com/MizrahiTeam/Educage). The Educage is an automated home-cage training apparatus designed to be used simultaneously with several mice. One advantage of the Educage over other procedures is that human interference is brought to minimum and training efficiency increases. The ‘Educage’ is a small modular chamber (10×10×10 cm), opening on one end into a standard animal home cage where mice can enter and exit freely (Fig. 1a). On its other end, the chamber contains the hardware that drives the system, hardware for identifying mice and measuring behavioral performance (Fig. 1a, Fig. S1a, and Methods). Mice were free to engage the behavioral port at their own will, where they consume all of their water intake. Following habituation, mice were trained on a go/no-go auditory discrimination task to lick in response to a target tone (a series of 10 kHz pure tones) and withhold licking in response to the non-target tone. A ‘lick’ response to a target was rewarded with a water drop (15 microliter) and considered as a ‘Hit’ trial. A ‘no lick’ response to a target sound was considered as a ‘Miss’ trial. A lick response to the non-target was considered a ‘False Alarm’ (FA) trial, which was negatively reinforced by a mild air puff (60 PSI; 600ms) followed by a 9 second ‘timeout’. A ‘no lick’ response to the non-target was considered a correct rejection (CR) and was not rewarded (Fig. 1a). On average, mice performed 327 ± 71 trials per day, mainly during dark hours (Fig. S1b).

The initial level of learning was to identify a non-target stimulus (a series of 7.1 kHz pure tones) from the 10 kHz target stimulus. These stimuli are separated by 49% of an octave and are perceptually easily separated by mice. Despite the simplicity of the task, behavioral performance varied widely between mice (Fig. 1b, Fig. S1c). On average, it took mice 54±38 trials to cross our criterion of learning, which was set arbitrarily at d’=1 (Fig. 1b; dotted line), and gradually increased to plateau at d’=2.35±0.64 (Fig. 1d). To extend the task to more challenging levels, we gradually increased task difficulty by changing the non-target tone closer to the target tone. The target tone remained constant at 10 kHz throughout the experiment and only the non-target stimulus changed. The lowest distance used between target and non-target was 3%/octave (9.6 kHz vs 10 kHz). A representative example from one mouse’s performance in the Educage throughout a complete experiment is shown in figure 1c. The just noticeable difference (JND) for each mouse was determined when mice could no longer discriminate (e.g. the JND of the mouse shown in figure 1c was determined between 6-10%/octave). The range of JND’s was 3-14%/octave and averaged 8.6±4.7%/octave. These values of JND are typical for frequency discrimination in mice ^31,49^. Most mice improved their performance with training (Fig. 1e, f), showing improved perceptual abilities along the task. The duration to reach JND varied as well, ranging 3069±1099 trials (14±3 days). Detection times, defined as the time in which lick patterns in the correct reject trials diverged from the lick patterns of the hit trials, increased monotonically by ~177 ms per each step of task difficulty demonstrating the increased perceptual load during the harder tasks (Fig. S1d).

To show that gradual training is necessary for perceptual learning ^5,8,11^ we trained groups of littermate mice on different protocols simultaneously. In one group of mice we used a standard protocol and the animals were trained on the gradually increasing task difficulty described above. Simultaneously, in the second group of mice – termed ‘easy only’ – animals were trained continuously on the easy task. Although both groups of mice trained together, only the mice that underwent gradual training were able to perform the hard task (Fig. 1g). Taking together, these data demonstrate the efficiency of the Educage to train groups of mice to become experts in discriminating between a narrowband of frequencies in a relatively short time and with minimal human intervention.

### Representation of pure tones in A1 following perceptual learning

To evaluate cortical plasticity following perceptual learning, we compared how pure tones are represented in A1 of expert mice, trained to discriminate between narrowband frequencies, and age matched naive mice who were never introduced to these tones. We used *in vivo* loose patch recording of L2/3 neurons in anesthetized mice to record tone-evoked spiking activity in response to 3-40kHz pure tones (Fig. 2a,b; Table 1). We targeted our recording electrode to the center of A1 based on previously validated stereotactic coordinates ^40^. Loose patch recording enables superb spatial resolution, is not biased to specific cell types and has superb signal to noise ratio for spike detection. However, one caveat of this technique is a potential bias of recording sites along the tonotopic axis. In order to overcome it we measured from neurons in a large number of animals, such that possible biases are averaged out. In naïve mice, responses were highly heterogeneous, with best frequencies covering the whole frequency range (Fig. 2c,d; n=105 neurons, n=22 mice, red). In expert mice, best frequencies of tuning curves were biased towards the frequencies that were presented during learning (Fig. 2c,d; n=107 neurons, n=21 mice, blue). These data show, as expected from previous literature, that learned frequencies in A1 become overrepresented at least as measured by the neuron’s best frequency (BF). We next showed that this overrepresentation was specific to the learned tone by training mice on 4 kHz as the target tone and recorded neurons in a similar manner to the abovementioned experiment. Indeed, L2/3 neurons in A1 of mice training on 4kHz showed BF shifts towards 4 kHz (Fig. S2a). Temporal responses to the trained frequencies were only slightly different between naive and expert mice. Specifically, average spiking responses were slightly but significantly faster and stronger in experts (Fig. S2b, Table 1).

**Figure 2-.**
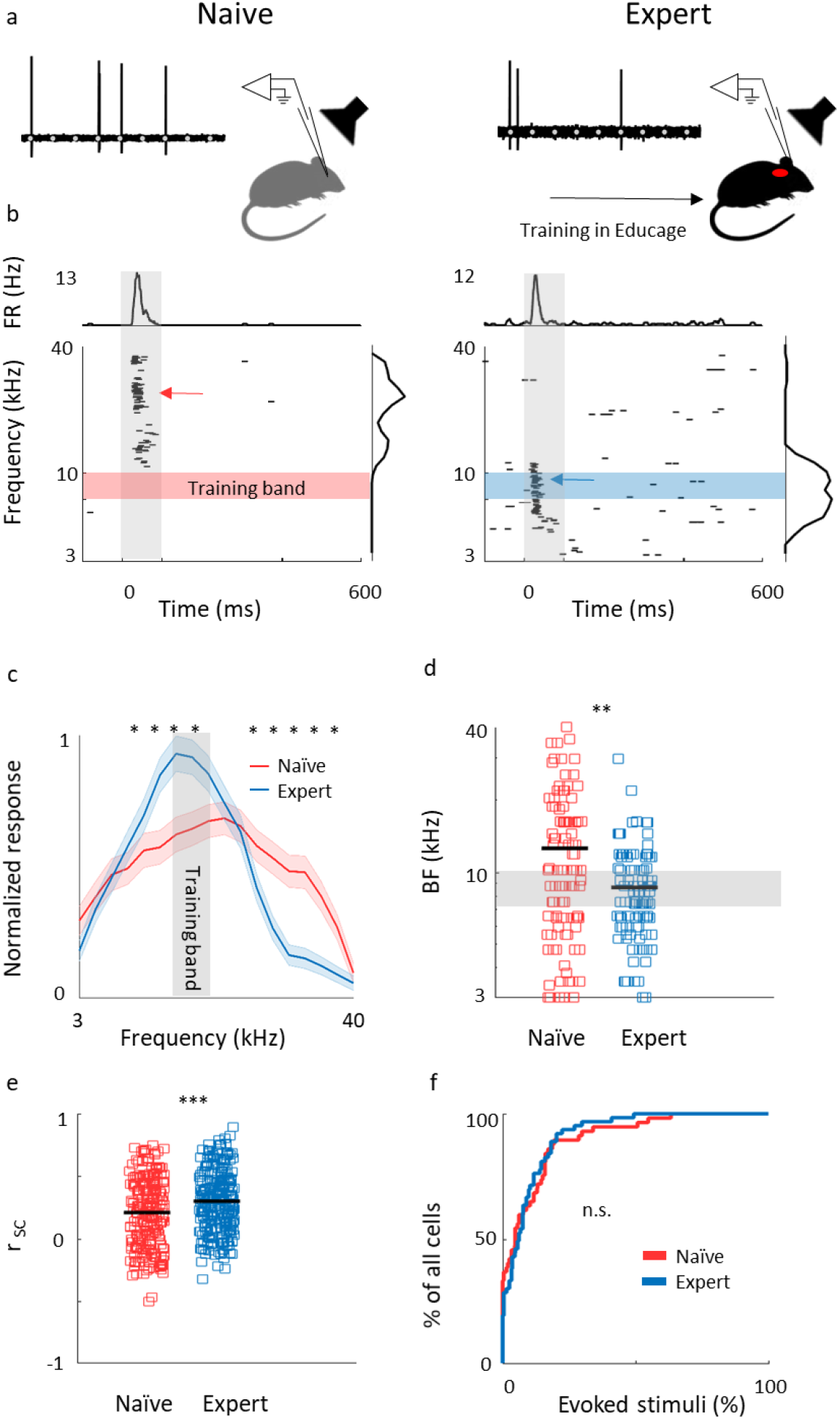
Learning induces over representation of the learned stimuli. **A.** Schematic representation of the experimental setup and a sample of the loose patch recording showing a representative cell from naïve (left) and expert (right) mice (gray markers indicate tone stimuli). **B.** Raster plots and peri-stimulus time histograms (PSTH) in response to pure tones of the cells shown. Gray bars indicate the time of stimulus presentation (100 ms). Color bars and arrows indicate the training frequency band (7.1-10 kHz) and BF, respectively. **C.** Population average of normalized response tuning curves of 105 neurons from naïve mice (red) and 107 neurons from expert mice (blue; mean ± s.e.m). Gray area indicates the training frequency band. Asterisks correspond to frequencies with significant response difference (Mann-Whitney U-test: *p<0.05). **D.** Best frequency (BF) of individual neurons from naïve mice (red markers) and expert mice (blue markers; Mann-Whitney U-test: **p<0.01). **E.** Pairwise signal correlations (rsc) values between all neighboring neuronal pairs in naïve (red) and expert (blue) mice. Neurons in expert mice have higher rsc (Mann-Whitney U-test: *** p<0.001). **F.** Cumulative distribution of response selectivity in naïve (red) and expert (blue) mice. Response selectivity was determined as the % of all frequency-intensity combinations that evoked a significant response. Distributions are not significantly different (Kolmogorov Smirnov test; p=0.542).

**Table 1-.** Learning induced physiological changes. A summary table of the complete dataset of recordings from excitatory neurons in naïve and expert mice after perceptual learning of pure tones. Columns show different parameters of the dataset or property tested. The third row shows the statistical p value between naïve and experts using a Mann-Whitney U-test.

| Group | Animals (n) | Cells (n) | BF (kHz) | Spontaneous spike rate (Hz) | Evoked firing rate (Hz) | FR in T.B (Hz) | Response latency (ms) | Selectivity (% of all stimuli) |
| --- | --- | --- | --- | --- | --- | --- | --- | --- |
| Naïve | 22 | 105 | 11.3 ± 8.4 | 0.45±0.58 | 15.5 ± 5.3 | 3.4 ± 5 | 35.2±8.7 | 9.7±13.9 |
| Expert | 21 | 107 | 8.6 ± 4 | 0.59±0.65 | 13.2 ± 3.4 | 4.2 ± 4 | 32.6±9.7 | 8.5±9.9 |
| Mann-Whitney U-test |  |  | 0.004 | 0.06 | 0.01 | 0.02 | 0.007 | 0.58 |

We recorded the responses to pure tones at different intensities and constructed frequency response areas (Fig. S2c). Pairwise signal correlation of neighboring neurons, calculated from these frequency response areas were high in naïve mice (0.2±0.28) but even higher in experts (0.3±0.24; Fig. 2E). Notably, the increased signal correlation was not an artifact of differences in response properties between naïve and expert group (Fig. S2d) but reflected true similarity in receptive fields (Fig. S2e). Thus, the basal level of functional heterogeneity in A1 ^40,50,51^ is reduced following learning. This learning-induced increase in functional homogeneity of the local circuit, emphasizes the kind of shift that local circuits undergo. Since neurons in expert mice did not have wider response areas (Fig. 2F, Table 1), our data suggests that neurons shifted their response properties towards the learned tones at the expense of frequencies outside the training band. These results are largely consistent with previous studies in monkeys, cats, and rats (reviewed in ^21^), extending this phenomenon of learning-induced changes in tuning curves to the mouse, to L2/3 neurons and to local circuits.

### A generative model of A1 population responses to pure tones

To what extent does overrepresentation of a learned stimulus sub-serve better discrimination by the neural population? To answer this question, we built a statistical model of tuning curves of neurons in A1 using six basis functions (Fig. S3), that correspond to the six largest Singular Value Decomposition (SVD) vectors of the population responses (Independent Basis Functions (IBF) method; see Methods). In figure 3a we show two representative examples of tuning curves and their reconstruction by our model. In contrast to Gaussian fits used previously^46,52^, our model captures the salient features of the shape of the auditory tuning curves (asymmetry, multimodality), yet also smoothed the *raw* response vectors to reduce overfitting due to finite sampling.

**Figure 3-.**
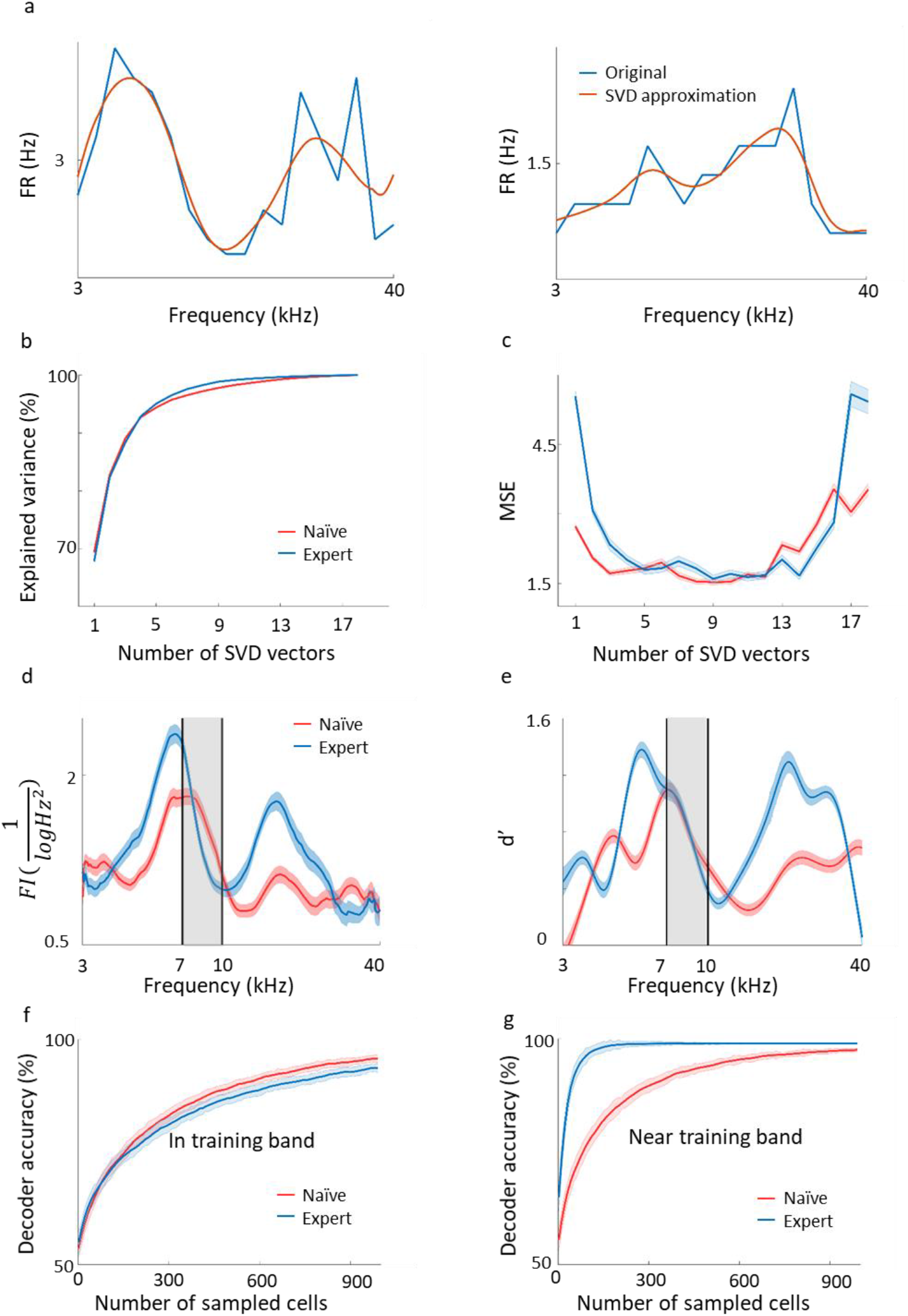
Plasticity in A1 does not improve the discrimination of the learned tones. **A.** Two representative examples of tuning curves of neurons recorded in A1. Average spike rates are shown in blue and the SVD approximation of the particular curves are shown in orange. Note that although the SVD approximation is smooth, it captures the irregular dynamics (i.e. non-Gaussian) of the tuning curves. **B,C.** The explained variance and error as a measure of the number of SVD vectors used in the model. **D.** Fisher Information calculated form the tuning curves of the both populations along the frequency dimension. Note the increased FI for the expert neurons in the flanks of the training band but not within it (gray band). **E.** Discriminability (d’) of SVM decoder along the range of frequencies. Pairwise comparison along the continuum are performed for frequencies 0.2198 octave apart. In accordance with ‘d’ the decoder does not perform better in the training band (gray shade). **F,G.** Classification performance of the decoder as a function of the number of neurons in the model. In the training band (F), the performance is similar for both naïve and expert mice. Outside the training band (0.4396 octave apart; G), performance improved rapidly in the expert mice.

In order to choose the appropriate number of basis functions, we determined the minimal number of basis functions that achieves good performance in reconstructing test single trial responses. In figure 3b we show the fraction of explained variance as a function of the number of basis functions*, K*. For both the naïve and expert groups the explained variance reaches above 96% after five basis functions. Figure 3c shows the Mean Square Error (MSE) on the unseen trials as a function of *K* in data from both the naïve and expert animals. The MSE exhibits a broad minimum for *K* in a range between 6-13. Interestingly, both naïve and experts achieve roughly the same MSE values, although for experts the MSE values at both low and large values of K are considerably larger than that of the naïve, presumably due to the smaller number of cells in the expert conditions. Taken together, we conclude that for these conditions six basis functions are the appropriate number and used this value for our calculations.

Based on the IBF method described above, we generated a population of tuning curves (500 ‘new neurons’), and estimated their total Fisher Information (FI; see Methods). Figure 3d shows the FI as a function of the stimulus *f* for both the naïve and expert conditions. Surprisingly, the FI of the neurons from expert animals was enhanced relative to the naïve group, but only for stimuli at both flanks of the training band. Importantly, the FI within the band of the trained frequencies remained unchanged (Fig. 3d, within the black lines). The same result holds true for a performance of a support vector machine (SVM) classifier. Using SVM to separate any two frequencies that are 0.2198 octave apart, discriminability (d’) values derived from the classifier’s error show similar results as the FI (Fig. 3e; ^45^). The value of d’ is larger in the expert groups as compared to the naïve but only outside the training band, whereas within the band discriminability is not improved (or even slightly compromised).

The results shown in figure 3 do not change qualitatively if we use our SVD model for the recorded neurons, as opposed to newly modelled neurons. One advantage of having a generative model for the population responses is that we can generate an unlimited number of trials and tuning curves. We took advantage of this to explore whether the results of the FI and SVM change with population size. To answer this question, we evaluated the mean discrimination performance (over test neurons) as a function of the number of sampled cells, *N*, which increases as expected. Consistent with the results of the SVM, the performances in the naïve and expert groups are similar with slight tendency for a higher accuracy in the naive population at large *Ns* (Fig. *3f)*. In contrast, for frequencies near the training band, the accuracy is substantially larger in the expert than in the naïve group for virtually all *N* (Fig. 3g). Thus, it seems that learning induced changes in tuning curves that do not improve discriminability of the learned stimuli.

### Perceptual learning of natural sounds

Natural sounds are characterized by rich spectro-temporal structures with frequency and amplitude modulations over time ^53^. Discrimination of such complex stimuli could be different from that of pure tones. Thus, we next designed a task similar to that with the pure tones but using mouse vocalizations as the training stimuli. We used playback of pups’ wriggling calls (WC) as the target stimulus (Fig. 4a - top). As the non-target stimuli, we used frequency modulations of the WC; a manipulation that allowed us to morph one stimulus to another by a continuous metric (Fig. 4a). The range of sound modulation used here was indexed as a “speeding factor” (see Methods for details). In short, a modulation factor of 0.66 affected the original WC more than a modulation factor of 0.9 did, and is therefore easier to discriminate (Fig. 4b). To reach perceptual limits we trained mice gradually, starting with an easy version of the task (WC vs a temporally reversed version of the WC) and then gradually to modulated calls starting at 0.66 modulation. Once mice reached >80% hit rates we changed the non-target stimulus to more difficult stimuli until mice could no longer discriminate (Fig. 4c). Mice (n=11) learned the easy task, i.e., discriminating WC from a 0.66 modulated call, with average d’ values of 2.5±0.4 (Fig.4d). On average, mice could only barely discriminate between a WC and its 0.9 modulation (d’ at 0.9 was 1±0.8; Fig. 4d). While these discrimination values were comparable to the performance of pure tones, detection times were substantially higher (Fig. S4a). For similar d’ values, discriminating between the vocalizations took 300-1000ms longer as compared to the pure tone tasks (Fig.4e). In addition, learning curves were slower for the vocalization task as compared to the pure tones task. The average number of trials to reach d’=1 for vocalizations was 195 trials, more than three times longer as compared with pure tones (compare Fig. S4b and 1b, respectively). These differences may arise from the difference in the delay, inter-syllable interval and temporal modulation of each stimulus type, which we did not further explore. Taken together, these behavioral results demonstrate a gradual increase in perceptual difficulty using a manipulation of a natural sound.

**Figure 4-.**
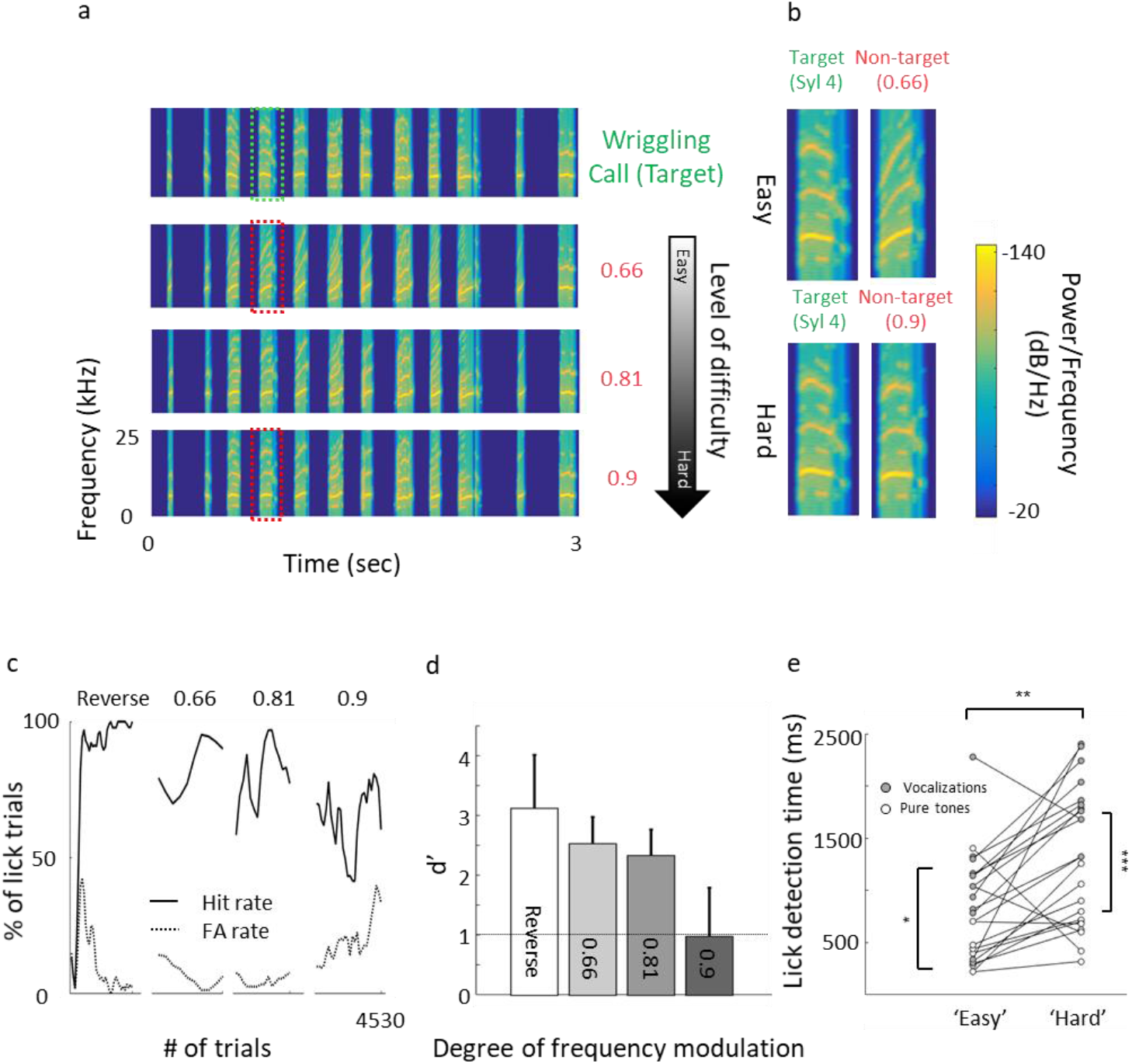
Perceptual learning of vocalizations. **A.** Spectrograms of the wriggling call (WC, ‘target’ stimulus, top panel) and the manipulated WC’s (‘non-target’ stimuli, bottom panels). **B.** Enlargement of the spectrogram’s 4^th^ syllable of the target WC and the manipulated calls. Top, a large manipulation (speeding factor-0.66), which is perceptually easy to discriminate from the WC. Bottom, a minor manipulation (speeding factor-0.9), which is perceptually closer to the WC. **C.** Lick responses to the target tone (solid line) and non-target tone (dashed line), binned over 50 trials, of one representative mouse along the different discrimination stages. The task of the first stage was to discriminate between WC vs reversed playback of the call (‘Reverse’). The following stages are different degrees of call modulation. Titles correspond to the speeding factors used for the non-target stimulus. **D.** Population average d’ values for the different discrimination levels. N=9 mice (mean ± s.e.m). Shades denote the level of difficulty. **E.** Comparison between detection times during the easy and difficult stages of pure tone (blank circles) and vocalizations (filled circles) discrimination tasks. Detection times are significantly different between all groups (Mann-Whitney U-test: ***p<0.001).

### Sparser response in A1 following perceptual learning of natural sounds

To study the neural correlates in A1 that support natural sound discrimination we recorded L2/3 neurons in response to the learned stimuli (Fig. 5a), expecting increased representation of these particular stimuli. Surprisingly, and in contrast to the results from pure tone discrimination, we did not find an increase in the representation of the learned stimuli. The fraction of cells responding to the trained vocalization remained constant (Fig. S5a) as well as the evoked firing rate for the preferred vocalization or the preferred syllable within a vocalization (Table 2).

**Figure 5-.**
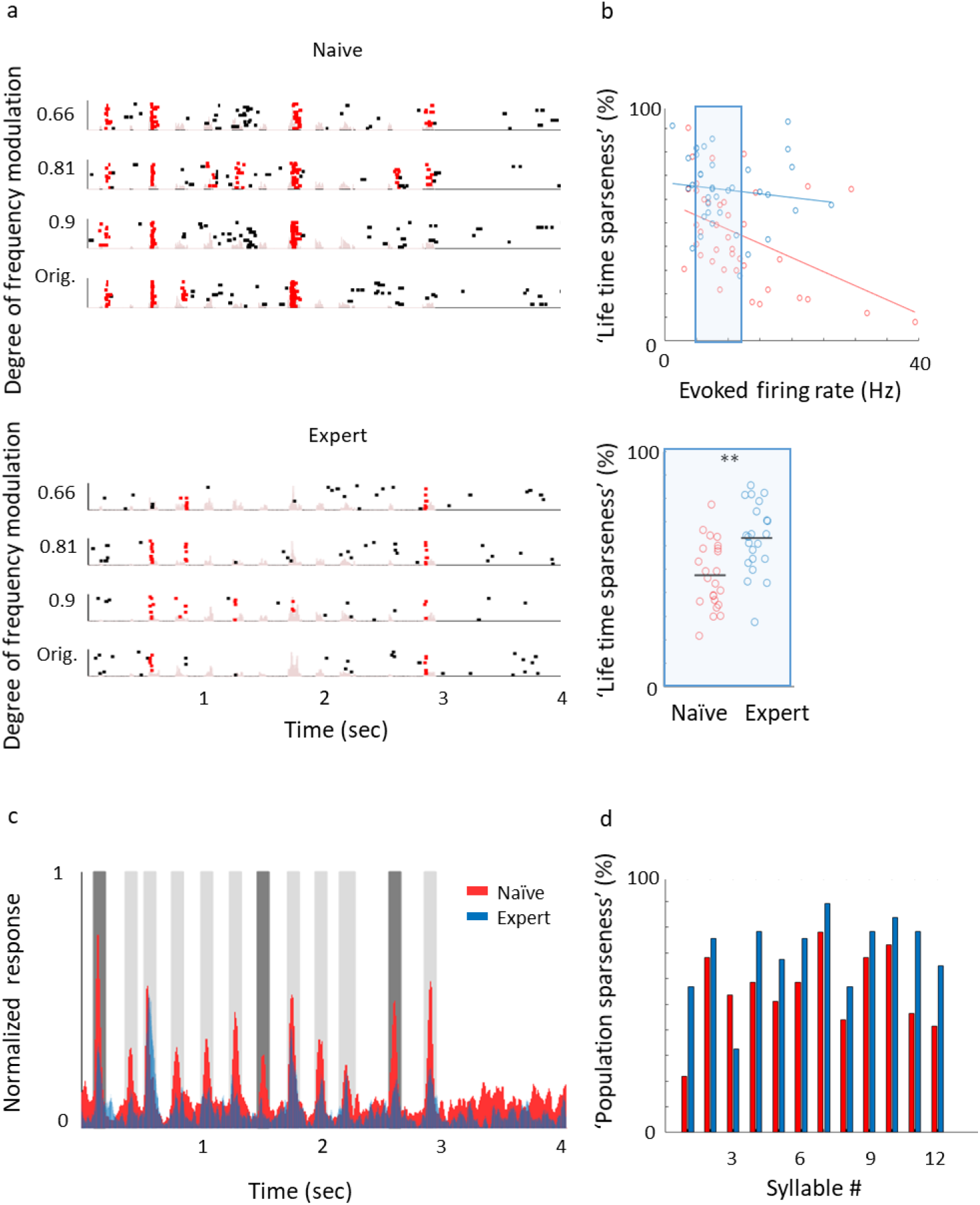
Learning complex sounds induces ‘sparsening’. **A.** Representative examples of raster plots in response to 4 modulated wriggling calls from naïve (top) and expert (bottom) mice. Red lines indicate spikes in response windows which are significantly above baseline. The stimulus power spectrum shown in light pink in the background. **B.** Top: Scatter plot of ‘life time sparseness’ (along the vocalization, see methods) and evoked firing rate for individual neurons from naïve (red circles) and expert (blue circles) mice. The middle range of firing rate distribution (0.5 SD bellow and above the median) is indicated as a blue rectangle. Bottom: Life time sparseness for all cells from the middle range of the distribution show significant difference between the groups (Mann-Whitney U-test; p=0.001). **C.** Average normalized PSTHs calculated from all neurons in response to the original WC. Data is shown overlaid for naïve (red) and expert (blue) mice. In expert mice, only syllables 1, 7 and 11 evoked significantly weaker responses as compared to naïve mice (dark gray bars, Mann-Whitney U-test; p=0.03, 0.001, 0.03). **D.** Population sparseness (% of cells that evoked a significant evoked response to a given syllable) of all neurons in naïve (red) and expert (blue) for the different syllables in the call.

**Table 2-.** Vocalization learning induced physiological changes. A summary table of the complete dataset of recordings from excitatory neurons in naïve and expert mice after perceptual learning of vocalizations. Columns show different parameters of the dataset or property tested. The third row shows the statistical p value between naïve and experts using a Mann-Whitney U-test.

| Group | Animals (n) | Cells (n) | Spontaneous firing rate (Hz) | Evoked firing rate (Hz) | Response latency (ms) | Response selectivity (% of evoked syllables) | Fano-Factor |
| --- | --- | --- | --- | --- | --- | --- | --- |
| Naïve | 8 | 41 | 1.8±2.9 | 7.6±5.9 | 36±11 | 25.4±21.2 | 1 ± 0.45 |
| Expert | 6 | 37 | 0.77±0.7 | 6.6±3.6 | 43±13 | 15.3 ± 12.2 | 1.1 ± 0.35 |
| Mann-Whitney U-test |  |  | P=0.1 | P=0.48 | P=0.02 | p=0.046 | P=0.12 |

Instead, representation in expert mice trained on vocalizations was sparser. Specifically, we measured the ‘lifetime sparseness’ ^47,48^ of each neuron and found that neurons of expert mice had higher sparseness and were more selective in their response within the call, regardless of their evoked firing rate (Fig. 5b; Naïve: 47±14%; Expert: 63±15%; p=0.001). Sparseness was also evident from the average population response to the vocalization (Fig. 5c). Nearly all syllables had weaker responses, three of which were statistically weaker (Fig. 5c, gray bars). Moreover, the population sparseness^48^ for almost all syllables in the call were higher in expert animal (Fig. 5d; Naïve: 55±16%; Expert: 70±15%; p=0.026). Notably, the smaller fraction of responses was not just an apparent sparseness due to increase in trial to trial variability, as reliability of responses by neurons in the expert group remained similar to reliability of responses in the naïve group (Table 2). Thus, ‘sparsening’ of A1 responses is a main feature of plasticity following learning to discriminate complex sounds.

We next asked whether the ‘sparsening’ described above bears more information to the learned stimulus. We analyzed population responses by including all neurons from all mice as if they are a single population (naïve: n=41 neurons from 8 mice; expert: n=37 neurons from 6 mice). We calculated the Pearson correlation of the population response to all responses in a pairwise manner (Fig. 6a). As compared to naïve mice, the absolute levels of correlations in expert mice were significantly lower for nearly all pairs of comparisons (Fig. 6a, asterisks). As expected, weaker modulations of the call and, hence, high similarity among stimuli, were expressed as higher correlations in the neuronal responses (Fig. 6b). The pairs of stimuli which mice successfully discriminated in the behavior (0.66 *vs*. the original WC and 0.81 *vs*. the original WC), had significantly lower correlation in the expert mice (Fig. 6b, rank sum test, p<0.05). Responses to the more similar stimuli that were near perceptual thresholds (i.e. 0.9 *vs*. the original WC) were lower in expert mice, but not significantly (Fig. 6b, rank sum test, p>0.05). This reduced Pearson correlation suggested that plasticity in A1 supports better discrimination among the learned natural stimuli. Indeed, a SVM decoder performed consistently better in expert mice, discriminating more accurately the original WC from the manipulated ones (Fig. 6c). As expected, the decoder performance monotonically increased when utilizing the responses to more syllables in the call. However, in the expert mice, performances reached a plateau already half way through the call, suggesting that neuronal responses to the late part of the call carried no additional information useful for discrimination. Similarly, the correlation of the population responses along the call, shows that responses were separated already following the first syllable, but that the lowest level of the correlation was in the 5^th^ - 7^th^ syllables range, which then rapidly recovered by the end of the call (Fig. S5c). These findings are also consistent with the behavioral performance of the mice as decisions are made within the first 1.5 seconds of the trial (or the first 7 syllables of the sentence). Specifically, the head of the mouse is often retracted by the time the late syllables are played (Fig. 4e). Taken together, ‘sparser’ responses improve neural discrimination of learned natural sounds.

**Figure 6-.**
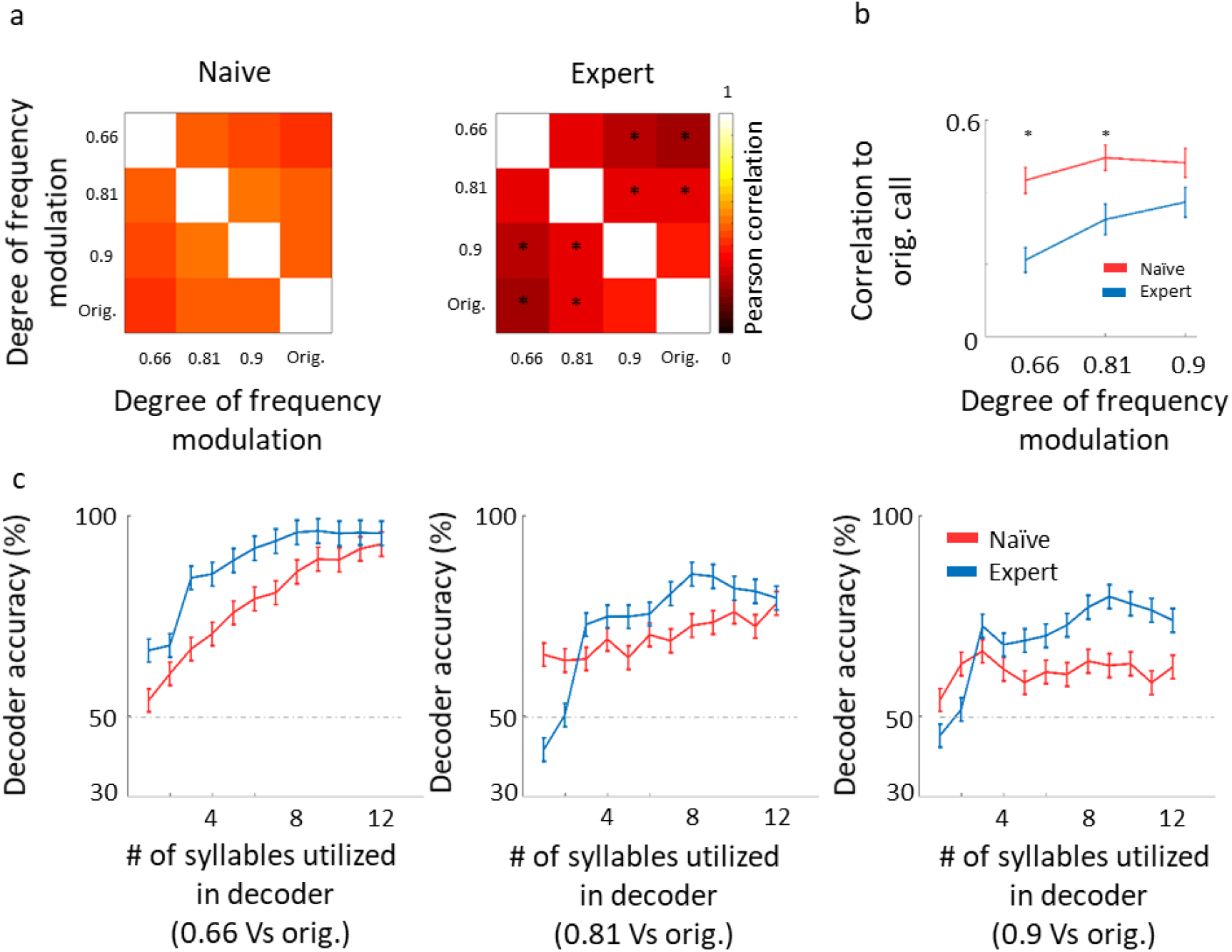
Decorrelation improves coding. **A.** Matrices describing the average response similarity of individual neurons between all combinations of stimuli in naïve (left) and expert (right) mice. Each pixel indicates the average Pearson correlation value calculated from all syllables’ evoked spike rate from all neurons to two different calls. Neurons from expert mice have lower correlation between responses to different modulated calls (asterisks indicate significant differences between naïve and expert groups; Mann-Whitney U-test, P<0.05). **B.** Pearson correlation between responses to modulated WCs and responses to the original WC in naïve (red) and expert (blue) mice (mean± s.e.m). Correlations are significantly different in the 0.66 and 0.81 modulation (Mann-Whitney U-test: * p<0.05). **C.** Classification performance of a Support Vector Machine (SVM) decoder. The decoder was tested for its accuracy to differentiate between the modulated WC stimuli against the original WC. The performance of the decoder is shown for neurons from the naïve (red) and expert (blue) groups. Decoder performance is plotted separately for the three different pairs of stimuli. Each point in each graph shows the number of syllables the decoder was trained on and allowed to use. Error bars are SEM for 1000 repetitions of leave-one-out cross validation.

### Learning induced plasticity of Parvalbumin neurons

The mechanisms responsible for the learning-induced changes following learning are currently unknown. We used mouse genetics and two-photon targeted patch to ask whether local inhibitory neurons could contribute to the observed plasticity we describe above. To this end, we focused only on parvalbumin inhibitory (PV^+^) interneurons as recent evidence indicates their role in a variety of learning related plasticity processes ^54–59^. Here we probe the role of inhibition in perceptual learning by measuring learning induce plasticity in the response properties of the PV^+^ neurons. We trained PV-Cre x Ai9 mice in the Educage and then patched single neurons under visual guidance (Fig. 7a; ^40,41^. In order to increase the sample of PV^+^ neurons we used targeted patch and often patched both PV^+^ and PV^-^ neurons in the same mice. PV^-^ neurons were used as proxy for excitatory neurons (these neurons were also included in the analysis shown in Fig. 2-6). All the TdTomato+ neurons that we patched were also verified as having a fast spike shape (Fig. 7a), a well-established electrophysiological signature of PV^+^ cells. PV^+^ neurons had response properties different from PV^-^ neurons in accordance with our previous work ^40^. For example, PV^+^ neurons responses were stronger and faster to both pure tones and natural sounds (Fig. 7b-c and Table 3; compare with Fig. 2b and 5a, see also ^40^).

**Figure 7-.**
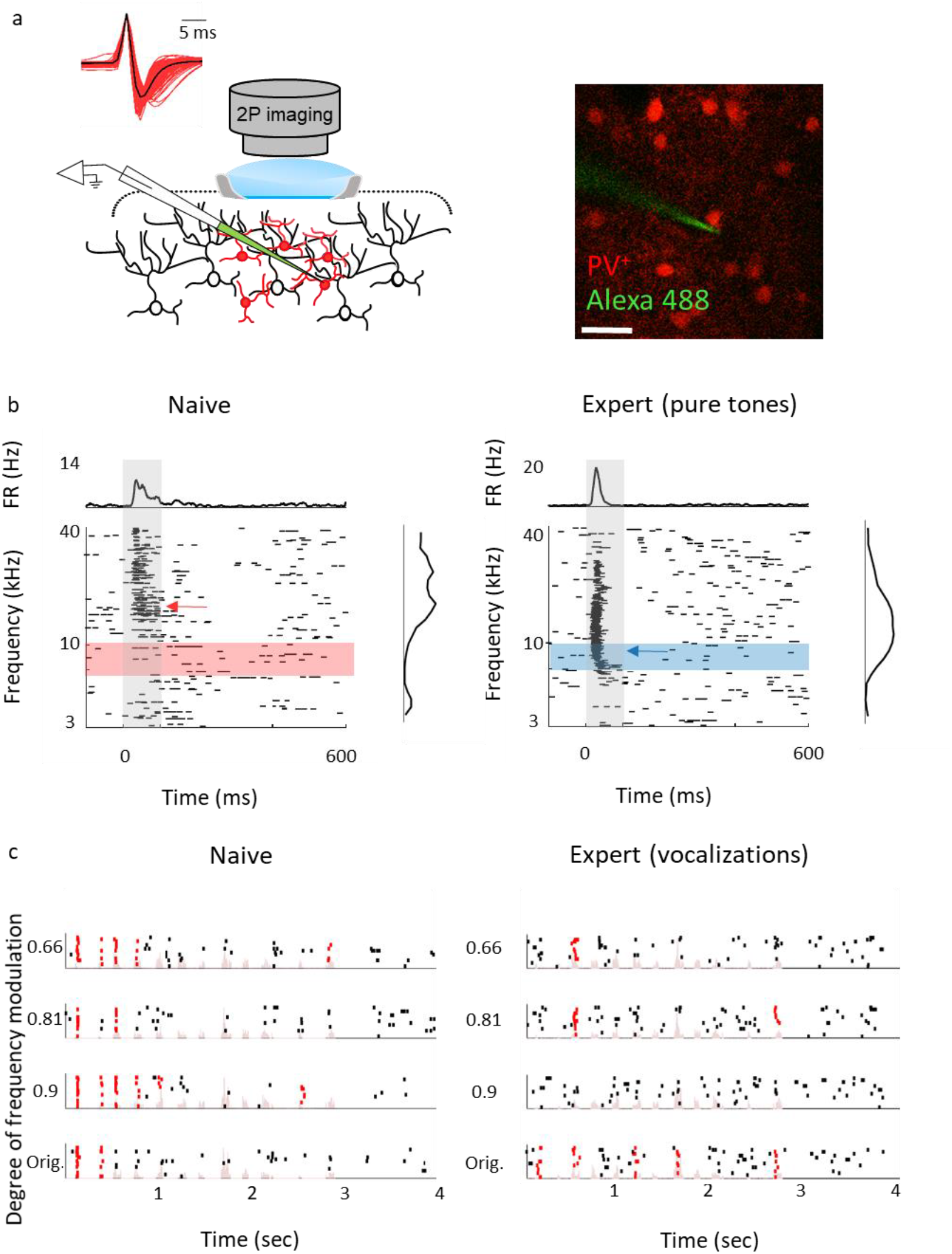
Response properties of PV^+^ neurons. **A. Left:** Schematic representation of the experimental setup for two-photon targeted patch and average spike waveform of 203 PV^+^ neurons. **Right:** Representative two-photon micrograph (projection image of 120 microns) of tdTomato+ cells (red) and the recording electrode (Alexa Fluor-488, green). **B.** Raster plots and peri-stimulus time histograms (PSTH) in response to pure tones of a representative PV^+^ neuron from naïve (left) and expert (right) mice. Gray bars indicate the time of stimulus presentation (100 ms). Color bars and arrows indicate the training frequency band (7.1-10 kHz) and BF, respectively. **C.** Representative examples of raster plots from PV^+^ neurons in response to 4 modulated wriggling calls from naïve (left) and expert (right) mice. Red lines indicate spikes in response windows which are significantly above baseline. Stimulus power spectrum shown in light pink in the background.

**Table 3-.**
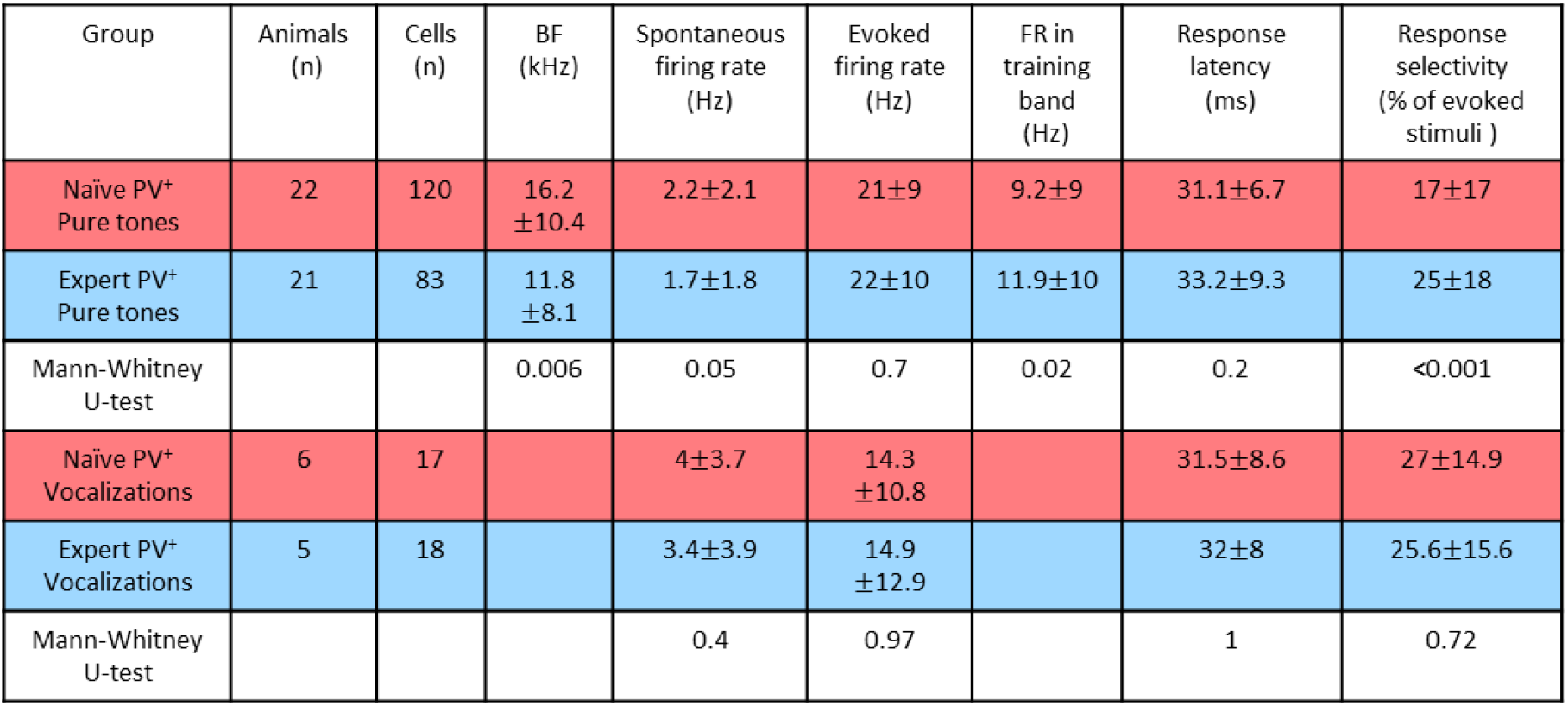
Physiological changes in PV^+^ neurons. A summary table of the complete dataset of recordings from PV neurons in naïve and expert mice after perceptual learning of pure tones (rows 1-2) and vocalizations (rows 4-5). Columns show different parameters of the dataset or property tested. The white color row shows the statistical p value between naïve and experts using a Mann-Whitney U-test for each group separately.

Following pure tone learning, PV^+^ neurons also changed their response profile. On average, the BF of PV^+^ neurons shifted towards the learned frequencies, similar to what we described for PV^-^ neurons (Fig. 8a,b). This result is consistent with recent evidence from the visual cortex showing increased selectivity to trained stimuli following learning ^60^.The shift in tuning curves of PV^+^ neurons was also accompanied by a significant widening of their receptive fields (Fig. S6a). When we compared the BF’s of PV^-^ and PV^+^ neurons within the same brain (within 250 microns of each other), we found that excitatory and inhibitory neurons became more functionally homogeneous as compared to naïve mice (Fig. 8c). As both neuronal groups show similar trends in the shift of their preferred frequencies we rule out a simple scenario whereas parvalbumin neurons increase their responses in the sidebands of the learning frequency. In other words, plasticity does not seem to be induced by lateral inhibition *via* parvalbumin neurons, but rather maintain a strict balance between excitation and inhibition, regardless of whether they are naïves or experts ^61,62^. Note that the peak of the PV^+^ population response and their BF distribution was on the outskirts of the training band, rather than within it (compare Fig. 8a,b with Fig. 2c,d), and concomitantly, the slope of the population responses at the trained frequency band increased due to learning (Fig. 8a). To assess the computational effect of the plasticity in the inhibitory neurons’ responses we have applied on their responses the same d’ and Fisher Information calculated for the PV^-^ neurons (Fig. 8d,e). Overall, the discrimination performance of the two cell populations (when equalized in size) is similar. However, the PV^+^ population shows a significant learning related increase in tone discrimination performance by these cells within the training band (Fig. 8d,e) in contrast to the results for PV^-^ neurons (Fig. 3d and e). This result is consistent with the above-mentioned increase in their response slopes in the trained frequency.

**Figure 8-.**
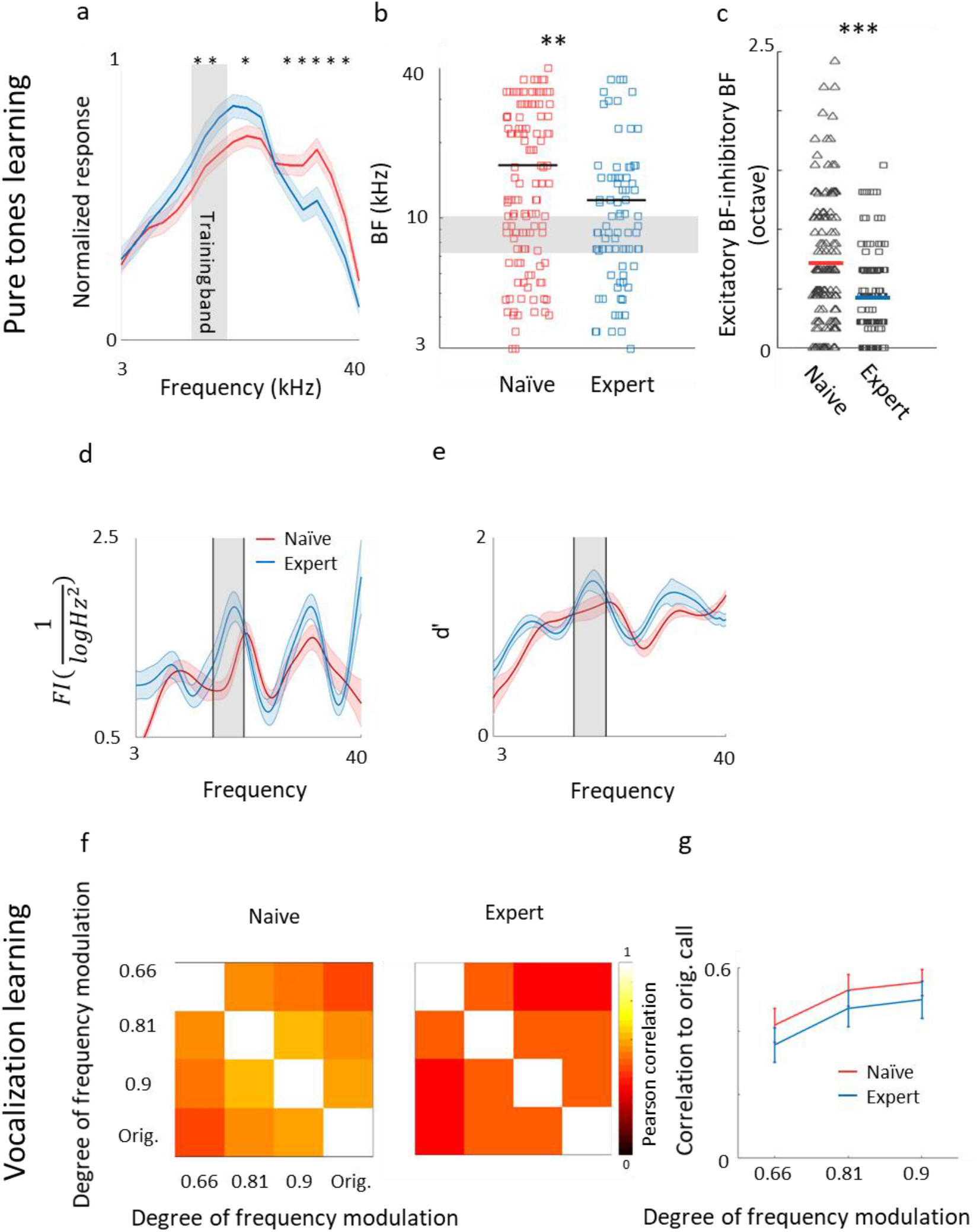
Plasticity of PV^+^ neurons after learning. **A.** Population average of normalized response tuning curves of n=120 PV^+^ neurons from naïve mice (red) and 83 PV^+^ neurons from expert mice (blue; mean ± s.e.m). Gray area indicates the training frequency band. Asterisks correspond to frequencies with significant response difference (Mann-Whitney U-test: *p<0.05). **B.** Best frequency (BF) of individual PV^+^ neurons from naïve mice (red) and expert mice (blue; Mann-Whitney U-test: **p<0.01) **C.** Average distance in BF between PV^-^ and PV^+^ neurons from the same penetration sites. Distances were significantly smaller in expert mice (Mann-Whitney U-test: ***p<0.001). **D.** Fisher Information calculated form the tuning curves of PV^+^ neurons **E.** Discriminability (d’) of SVM decoder along the range of frequencies. **F.** Matrices describing the average response similarity of individual PV^+^ neurons between all combinations of different stimuli in naïve (left) and expert (middle) mice. Each pixel indicates the average Pearson correlation value calculated from all syllables evoked spike rate from all neurons to two different calls. There was no significant difference between correlations of responses to different modulated calls in naïve and expert mice (Mann-Whitney U-test, P>0.05). **G.** Pearson correlation between responses to modulated WCs versus the responses to the original WCs in naïve (red) and expert (blue) mice (mean± s.e.m).). Correlations are not significantly different for all comparisons (Mann-Whitney U-test: p=0.4, 0.5, 0.5).

Following natural sound learning, we found no significant changes in basic response properties of the PV^+^ neurons (Table 3), nor in the degree of sparseness of their representation of the learned vocalizations (Fig. S6b). The relationship between the responses of PV^+^ and their PV^-^ neighbors remained constant as reflected in the similar slopes of the functions describing PV^-^ firing versus PV^+^ firing (Fig. S6c). This result suggests that the excitation-inhibition balance, as reflected in the responses of PV^-^ versus PV^+^, remains. In PV^+^ neurons, the temporal correlation along the call as well as the decoding performance from these neurons showed changes that are qualitatively similar to their PV^-^ counterparts, but the data across the population was noisier (Fig. 8f-g; Fig. S6d) perhaps due to the smaller sample of the PV^+^ dataset.

## DISCUSSION

### Plasticity in frequency tuning following perceptual learning

Shifts in the average stimulus representation towards the learned stimuli is not a new phenomenon. Similar findings were observed in numerous studies, multiple brain areas, animal models and sensory systems, including in auditory cortex ^10,16,63^. In fact, the model of learning-induced plasticity in A1, also known as tonotopic map expansion, is an exemplar in neuroscience ^18–20,64^; but see ^22,65–67^. Indeed, we also found that the number of neurons tuned to the learned frequency band increased in expert mice after perceptual learning (Fig. 2). Thus, our results support the observations of tuning curve plasticity in a primary sensory cortex, showing this here specifically for L2/3 neurons and extending it to A1 in mice.

Since we sampled only a small number of neurons in areas smaller than 250μm^3^, our observations cannot be inferred as direct evidence for tonotopic map expansion. Rather, our data emphasizes that plastic-shifts occur in local circuits (Fig. 2e). Given that neurons in A1 are functionally heterogeneous within local circuits ^40,50,51^, any area in A1 that represents a range of frequencies prior to learning could become more frequency-tuned once learned. Such a mechanism allows a wide range of modifications within local circuits to enable increased representation of the learned stimuli without necessarily perturbing gross tonotopic order. One advantage of local circuit heterogeneity is that it allows circuits to maintain a dynamic balance between plasticity and stability ^68^. Only mapping the full extent of auditory cortex, at both global and local scale and preferably over time in the same neurons ^69^, will enable to reveal the precise type of changes the cortex undergoes and how it exploits its variability to sub-serve perceptual learning.

Quantifying overrepresentation across a population of neurons and interpreting these in the context of the learned perceptual tasks have been the subject of extensive research ^70^. It has commonly been assumed that the learning-induced changes in tuning properties improves the accuracy of coding of the trained stimuli. In particular, for discrimination tasks between two stimuli differing by a small change in a one-dimensional stimulus parameter, perceptual learning theory predicts that sharpening of the slope of the tuning curves with respect to this parameter improves the discriminability of the stimuli ^45^. However, the observed increased representation of the BFs towards the training band in expert mice may not increase over all tuning slopes and may even decrease them, especially since the slopes tend to be small at the best frequencies. A closer look at the tuning curves in A1 shows that they are often irregular with multiple slopes and peaks (i.e. not having simple unimodal Gaussian shapes). Furthermore, the learning induced changes in the ensemble of tuning curves are not limited to shifting the BF, hence a more quantitative approach was required to assess the consequences of the observed learning induced plasticity on discrimination accuracy. This motivated us to develop a method that takes into account the irregularities in tuning curves of A1 neurons. To this end, we have used a new, SVD based, generative model (Fig. 3), allowing us to assess the combined effects of changes in BFs as well other changes in the shapes of the tuning curves. Surprisingly, both Fisher Information analysis as well as estimated classification errors of an optimal linear classifier, show that learning induced changes in tuning curves does not improve tone discriminability at trained values. This conclusion is consistent with previous work on the effect of exposure to tones during development. Early life experience during development which induces similar changes to those observed during learning has been argued to decrease tone discriminability for similar reasons ^71^. However, in that work, the functional effect of tuning curve changes was consistent with an observed impaired behavioral performance, suggesting that plasticity in A1 sub-serves discrimination behavior. In contrast, the stable (or even reduced) accuracy in the coding of the trained frequency we observed occurs despite the improved behavioral performance after training. What may this finding underlie?

The changes that we recorded in A1 following pure tone learning do not improve their contribution to discrimination of the task stimuli, and are thus orthogonal to direct behavioral performance. One interpretation of this finding is that learning induced plasticity in other brain regions, either in downstream (or parallel) auditory areas or in task related ‘readout’ areas enable improved behavioral performance. Notably, even the representation in naïve mice would be sufficient for reliable decoding at behavioral resolution – utilizing a population of only two-three hundred A1 neurons (Fig. 3g). Thus, changes in the tuning of A1 neurons are not necessary for discriminating the pure-tone task. Further, we suggest that the observed learning-induced changes in the tuning of pure tones in A1 are the result of unsupervised Hebbian learning induced by over exposure to the trained tones during the training period; similar to the reported results in early overexposure ^71^. Unsupervised learning signals are not driven by task related reward *per se* and may increase representation rather than discriminability. Increased representation of trained stimuli may lead to improved discriminability to untrained tones, as observed experimentally. Importantly, we find that for natural sounds, learning-induced changes in A1 responses improved the discriminability of the trained stimuli relative to the naïve responses. This suggests that A1’s primary function is in processing and coding complex auditory stimuli like e.g. natural sounds, rather than pure tones.

### Coding of Natural sounds in A1

Unlike pure tones, natural sounds are composed of rich spectro-temporal energies and include a set of sound features such as amplitude modulations, frequency modulations, harmonics, and noise. Although auditory cortex is organized tonotopically, it may not be essential for processing simple sounds, as these stimuli are accurately represented in earlier stages in the auditory hierarchy, as early as the brainstem ^53,72^. This is consistent with experiments showing that auditory cortex is necessary for associative fear conditioning with complex sounds ^54^ but not with pure tones ^73–75^.

A multitude of studies have shown that single neurons and populations in A1 respond to sounds in a non-linear fashion ^76–78^. For example, neurons in A1 can be selective to harmonic content that are prevalent in vocalizations and other natural stimuli ^79^. Some neurons in A1 show strong correlations to global stimulus statistics ^80^. Furthermore, neurons in A1 are sensitive to the fine-grained spectrotemporal environments of the sounds, expressed as strong gain modulation to local sound statistics ^81^, as well as to sound contrast and noise ^82,83^. All of these features (harmonics, globally and locally rich statistics, and noise) are well represented in the wriggling calls we played during learning. Which of these particular sensitivities changes after perceptual learning is not yet known but one expression of this plasticity can be the increased sparse sound representation we found here (Fig. 5, S5). Sparseness can take different forms ^84^. Here, sparseness, expressed both in the temporal pattern of individual neurons and at the level of the population, is mainly a result of reduced number of neurons in the network that respond to any of the 12 syllables played (Fig. S5). Such increase in selectivity to the syllables could arise from disparate mechanisms, and changes in the structure of local inhibition was one suspect that we tested ^85^.

### Inhibitory plasticity follows excitatory plasticity

Cortical inhibitory neurons are central players in many forms of learning ^86–88^. Inhibitory interneurons have been implicated as important for experience dependent plasticity in the developing auditory system ^89^, during fear learning in adulthood ^54,90^, and following injury ^91^. Surprisingly, however, and despite the numerous studies on parvalbumin neurons, we could not find any references in the literature of recordings from parvalbumin neurons after auditory perceptual learning. Two simple (non mutually exclusive) hypotheses are naively expected. One is that plasticity in inhibitory neurons are negative mirrors of the plasticity in excitatory neurons. This would predict that inhibitory neurons would increase their responses to the stimuli for which responses of excitatory neurons are downregulated (as found in the plasticity of somatostatin expressing (SOM) neurons following passive sound exposure (Kato et al. 2015), or in multisensory plasticity in mothers (Cohen and Mizrahi, 2015). The second is that inhibitory neurons enhance their response to ‘lateral’ stimuli thus enhancing selectivity to the trained stimulus, as suggested by the pattern of maternal related plasticity to pup calls (Galindo-Leon, Lin, and Liu 2009). To the best of our knowledge, our study thus provides a first test of these hypotheses in the context of perceptual learning. We found no evidence for these scenarios. Instead, a common motif in the local circuit was that parvalbumin neurons changed in a similar manner to their excitatory counterparts. These results are in line with the observation that PV neurons in V1 following visual discrimination task become as selective as their Pyramidal neighbors ^60^.

The cortex hosts several types of inhibitory cells ^92,93^, presumably serving distinct roles. While PV neurons are considered a rather homogeneous pool of neurons based on molecular signature, their role in coding sounds is not. This is evident in recordings from pyramidal neurons in A1 while optogenetically inhibiting the PV^+^ neurons resulted in mixed effects ^94,95^. In the visual cortex, PV^+^ cells’ activity (and presumably its plasticity) was correlated with stimulus-specific response potentiation but not in ocular dominance plasticity ^56^, again, suggesting that PV^+^ neurons are not necessarily involved in all forms of experience-dependent plasticity. Inhibitory neurons have been suggested to play a key role in enhancing the detection of behaviorally significant vocalization by lateral inhibition ^96^. But recent imaging data argue that somatostatin interneurons rather than PV^+^ interneurons govern lateral inhibition in A1 ^97^. Our results are also consistent with the observation that in contrast to the SOM neurons, the changes in responses following sound exposure are similar in PV^+^ and pyramidal neurons ^60,67^. To what extent somatostatin or other interneurons subtypes contribute to excitatory plasticity after auditory perceptual learning remains to be studied.

### Simplifying training with the Educage

Training rodents on perceptual learning paradigms has traditionally been performed manually over many weeks, sometimes months, of training ^11,18^. Behavior is confounded by all sorts of variability, even in the controlled conditions of a laboratory and within inbred mice of the same age and sex ^98^. The Educage is a new tool for standardizing cognitive tasks, as we demonstrated here for perceptual learning. The Educage allows training of larger animal datasets, effortlessly collecting thousands of trials per day, and effectively increasing statistical power for assessing inter-animal variability. Since mice train simultaneously, it potentially reduces stress-associated with factors like social deprivation. Other behavioral tools have been introduced in recent years ^31,32,99,100^. Each system has its own strengths or weaknesses in cost, design and operation load ^33^(e.g. see a new system for olfactory perceptual learning ^30^, and new systems with options for combining physiology and imaging ^28^). The simple design and operation of the ‘Educage’ together with its affordable hardware and software makes it yet another valuable tool for simplifying training.

## SUPPLEMENTARY FIGURES

**Supplementary figure 1.**
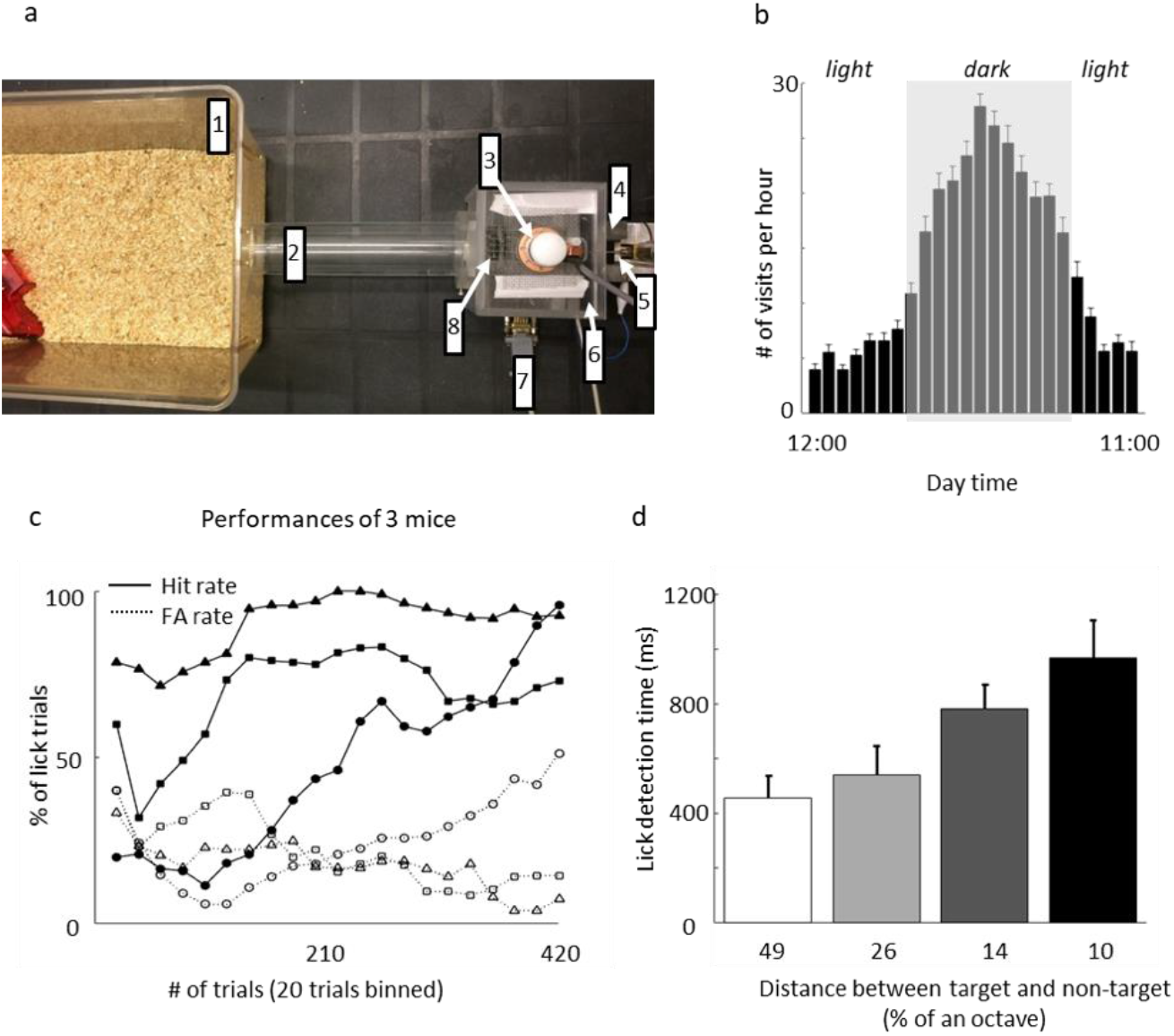
**A.** Photograph of the Educage platform (top view). 1. Home cage; 2. Connecting tunnel; 3. Auditory speaker; 4. IR diodes monitoring the behavioral port; 5. Water spout and licko-meter; 6. RFID antenna; 7. Connection to D/A converter; 8. Electrically conductive grid floor (for lickometer circuit). **B.** Histogram of daily hourly distribution of voluntary training. Gray area represents dark hours. N=39 mice (mean ± s.e.m). **C.** Representative examples of the performance of three mice (the same mice shown in ‘Figure 1B’) during the first stage of discrimination. The graph shows % of lick responses to the target tone (‘Hit rate’-solid line) and to the non-target tone (‘FA rate’-dashed line). **D.** Population averages of detection times for the different discrimination levels. N=12 mice (mean ± s.e.m). Shades denote the level of difficulty (from 49% to 10% octave apart).

**Supplementary figure 2.**
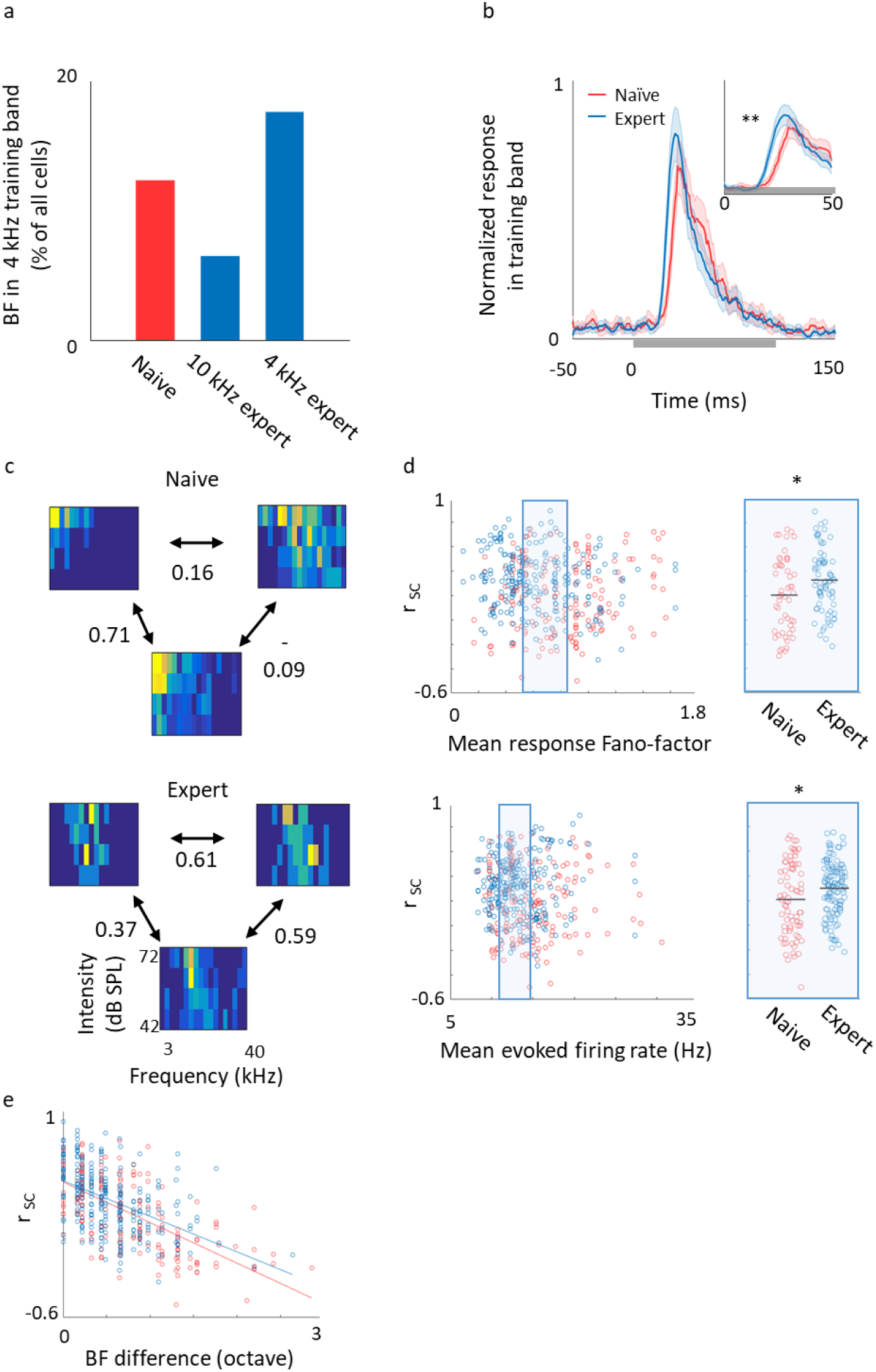
**A.** The fraction of neurons with BF that resides within a 4kHz frequency band in three different experimental groups. Naïve, (n=22 mice) - red; mice trained on a 10kHz (n=21 mice), and 4kHz (n=4 mice) as the target tone - blue. **B.** Population average of normalized PSTHs from naïve (red) and expert mice trained (blue; mean ± s.e.m). The PSTHs shown were collapsed for all intensities and for all frequencies inside the training band, binned at 1 ms time bins. Inset: zoom-in on the first 50 ms from stimulus onset. **C.** Representative examples of frequency response areas (FRA’s) and pairwise signal correlations (r_sc_) of neighboring neurons from the same local circuits in naïve (top) and expert (bottom) mice. **D.** Left, Scatter plot of pairwise r_sc_ and pair average of evoked firing rate or response Fano-factor for naive (red circles) and expert (blue circles). The range common to the naïve and expert (0.5 SD bellow and above the median) is indicated as blue rectangle. Right, All pairwise rsc values from the common range show consistent differences between groups, independent of these parameters (Mann-Whitney U-test: *** p<0.05). **E.** Scatter plot of pairwise rsc and pair best frequency difference showing strong dependency (Naïve: slope=-0.3; p=2E^-28^; R^2^= 0.42; Expert: slope=-0.27; p=5.8E^-17^; R^2^=0.22).

**Supplementary figure 3.**
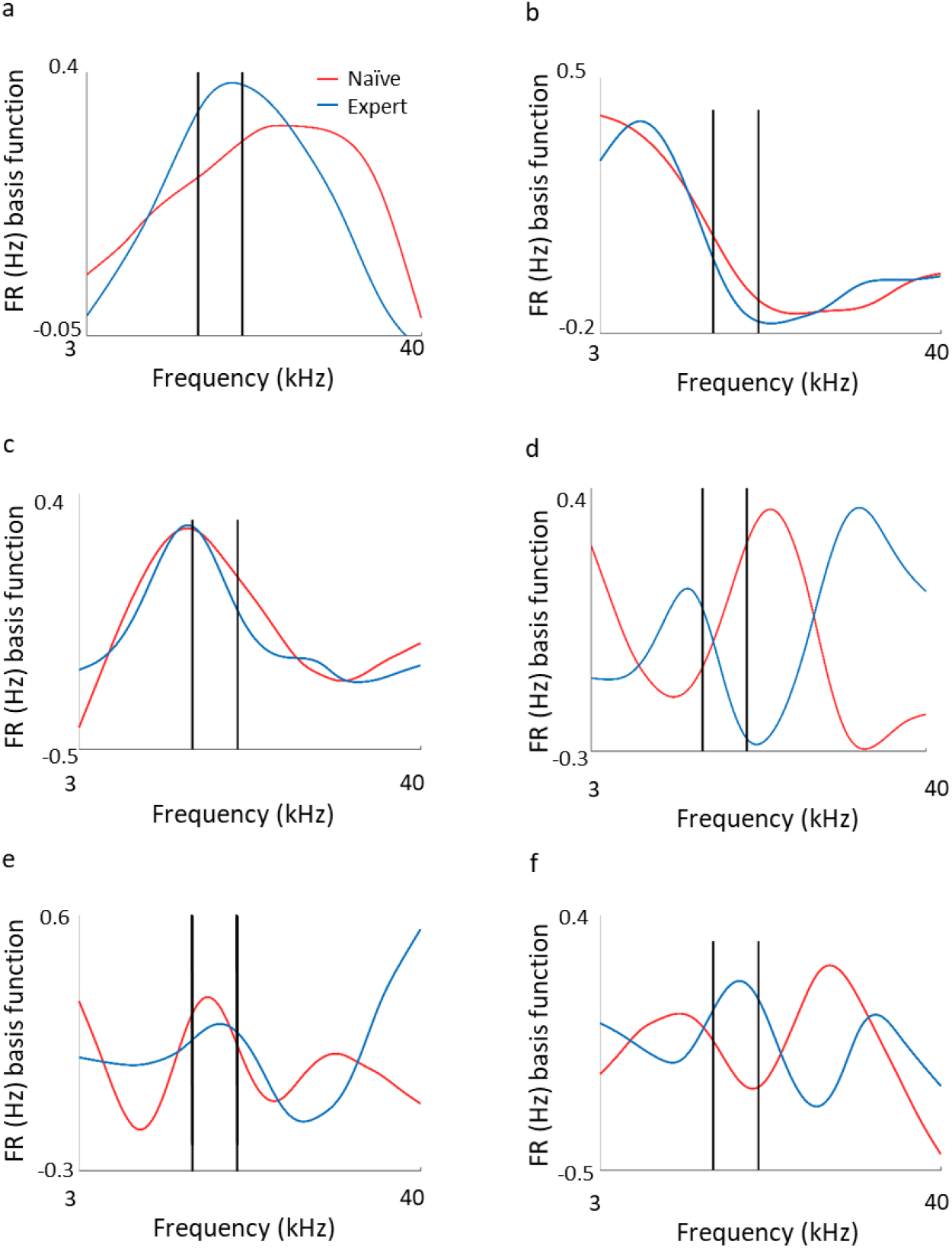
**A-F.** 6 basis functions coming out of the SVD analysis for naïve (red) and expert (blue) neurons

**Supplementary figure 4.**
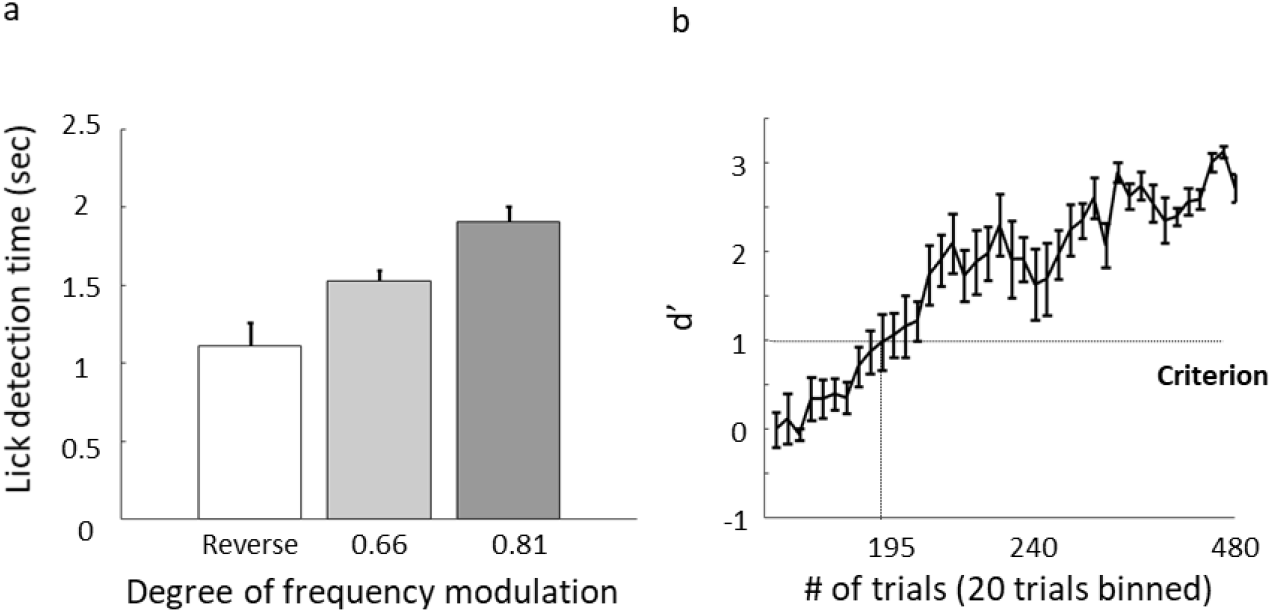
**A.** Population average of detection times for the different discrimination stages. N=9 mice (mean ± s.e.m). Shades denote the level of difficulty. **B.** d’ values (mean± s.e.m) for the first stage of discrimination (WC vs Reverse). The learning criterion is represented as dashed line (d’=1).

**Supplementary figure 5.**
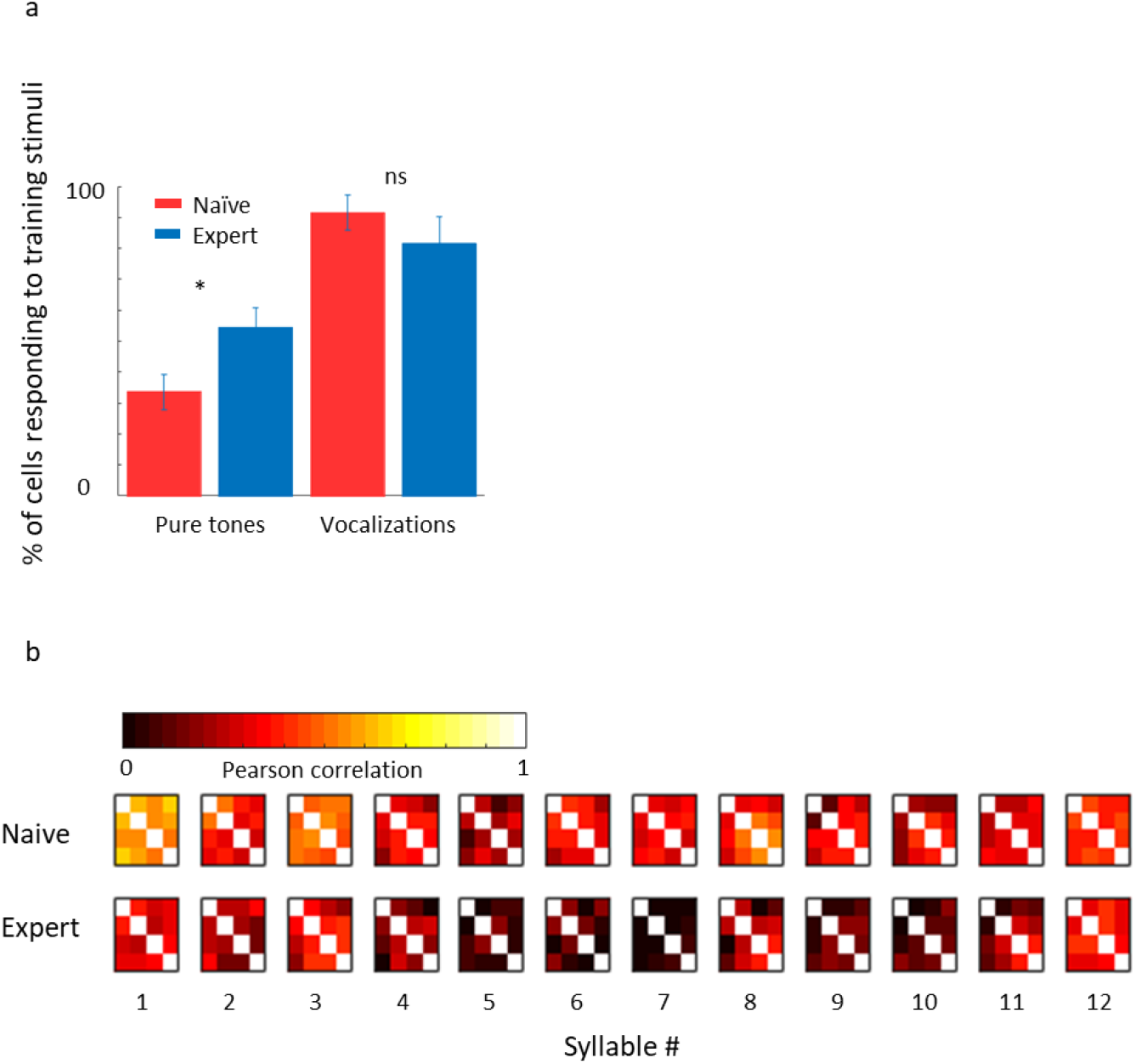
**A.** Fraction of cells with significant response to frequencies within the training band (left) and to the original WC in naïve (red) and expert (blue) mice (mean ± s.e.m). Fraction of cells responding to the trained vocalization remained constant (Mann-Whitney U-test: p=0.09) **B.** Correlation matrices for different syllables individually. Each pixel indicates the average Pearson correlation value from two different calls calculated from the evoked spike rate from all neurons in response to one syllable. Neurons from expert mice have lower correlation between responses to different modulated calls, and specifically in syllables 5-7.

**Supplementary figure 6.**
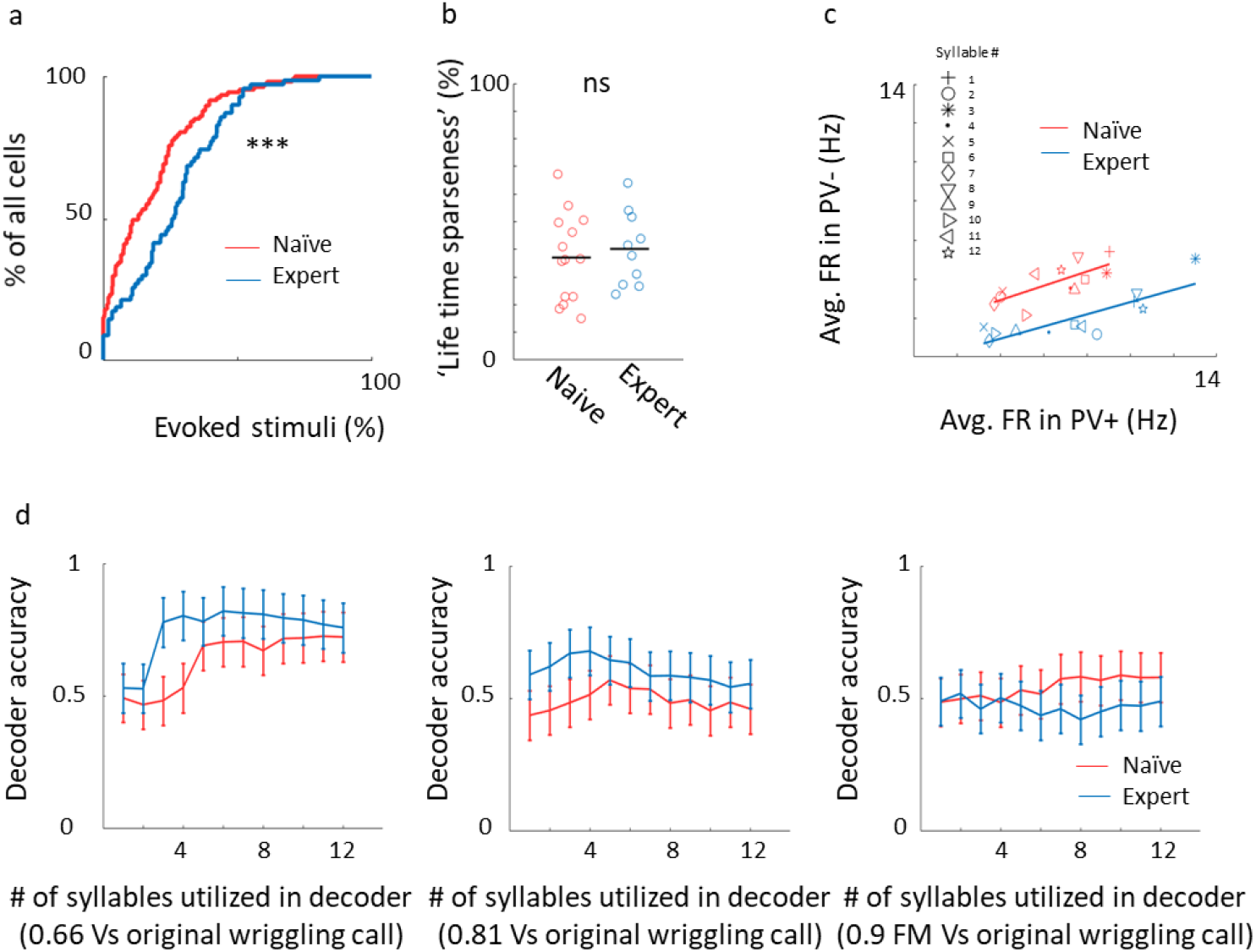
**A.** Cumulative distribution of response selectivity of PV^+^ neurons in naïve (red) and expert (blue) mice. Response selectivity was determined as the % of all frequency-intensity combinations that evoked significant response. PV^+^ neurons in expert were less selective (Kolmogorov Smirnov test; p=0.0015). **B.** Life time sparseness for all cells from the middle range of the firing rate distribution show no significant difference between naïve (red circles) and expert (blue circles) groups (Mann-Whitney U-test; p=0.46). **C.** Scatter plot showing linear dependency of average syllable evoked firing rate of PV^-^ and PV^+^ neurons in naïve (red; R^2^ = 0.5; slope=0.37; p=0.008) and expert (blue red; R^2^ = 0.7; slope=0.31; p<0.001) mice. Each marker corresponds to the average response for one syllable in the sentence. D. Classification performance of a Support Vector Machine (SVM) decoder based on PV^+^ responses.

## Acknowledgements

We thank Israel Nelken and Livia de Hoz for help in setting up the initial behavioral experiments. We thank Yoni Cohen, David Pash and Joseph Jubran for writing the software code of the Educage. We thank Yonatan Loewenstein, Merav Ahissar, and the Mizrahi lab members for comments on the manuscript. This work was supported by an ERC consolidators grant to A.M (#616063) and by Gatsby Charitable Foundation.

Author contributions
I.M. and A.M. designed the experiments. R.S.Z and H.S analyzed data. L.F. conducted experiments. Y.E. constructed the vocalization stimuli and software for their delivery. I.M. conducted experiments and analyzed the data. I.M. H.S. and A.M. wrote the paper.

